# Rising insecticide potency outweighs falling application rate to make US farmland increasingly hazardous to insects

**DOI:** 10.1101/715763

**Authors:** Margaret R. Douglas, Douglas B. Sponsler, Eric V. Lonsdorf, Christina M. Grozinger

**Affiliations:** Department of Environmental Studies & Environmental Science, Dickinson College, Carlisle, PA 17013, USA; Department of Entomology, Center for Pollinator Research, Huck Institutes of the Life Sciences, Pennsylvania State University, University Park 16802, PA, USA; Institute on the Environment, University of Minnesota, St Paul, MN 55108, USA

**Keywords:** pesticide, neonicotinoid, insect conservation, pest management

## Abstract

Each year, millions of kilograms of insecticides are applied to crops in the US. While insecticide use supports food, fuel, and fiber production, it can also threaten non-target organisms, a concern underscored by mounting evidence of widespread insect decline. Nevertheless, answers to basic questions about the spatiotemporal patterns of insecticide use remain elusive, due in part to the inherent complexity of insecticide use, and exacerbated by the dispersed nature of the relevant data, divided between several government repositories. Here, we integrate these public datasets to generate county-level annual estimates of total ‘insect toxic load’ (honey bee lethal doses) for insecticides applied in the US between 1997-2012, calculated separately for oral and contact toxicity. To explore the underlying drivers of the observed changes, we divide insect toxic load into the components of extent (area treated) and intensity (application rate x potency). We show that while contact-based insect toxic load remained relatively steady over the period of our analysis, oral-based insect toxic load increased roughly 9-fold, with reductions in application rate outweighed by disproportionate increases in potency (toxicity/kg) and increases in extent. This pattern varied markedly by region, with the greatest increases seen in Heartland and Northern Great Plains regions, likely driven by use of neonicotinoid seed treatments in corn and soybean. In this “potency paradox,” US farmland has become more hazardous to insects despite lower volumes of insecticides applied, raising serious concerns about insect conservation and highlighting the importance of integrative approaches to pesticide use monitoring.

**Significance statement:** Previous analyses disagree about whether US insecticide use is increasing or decreasing, a question of significant importance given the putative role of insecticides in recent insect declines. We integrated information from multiple national databases to estimate ‘insect toxic load’ (represented as honey bee lethal doses) of the agricultural insecticides applied in each US county from 1997 to 2012, and factors responsible for its change. Across the US, insect toxic load – calculated on the basis of oral toxicity – increased 9-fold. This increase was due to increases in the potency (toxicity/kg) of insecticides applied and in the area treated; the volume of insecticides applied declined. Toxic load increased most dramatically in regions where neonicotinoid seed treatments for field crops are commonly used.

## Introduction

Insects are the most diverse and abundant class of animals on earth, with an estimated 5.5 million species that dominate animal biomass in many ecosystems (1, 2). Given their ubiquity, it is not surprising that insect populations serve in key roles as both friend and foe to human societies. This is particularly true in agriculture, where farmers seek to manage populations of insect pests to produce essential food, fuel and fiber, a task in which insecticides play an important role. In the United States, agriculture accounts for ∼57% of insecticide weight applied and ∼85% of area treated, and so constitutes the single largest contributor to insecticide use (3, 4). However, since at least the 1960’s it has been widely recognized that insecticide application can also negatively affect non-target species, including populations of insect pollinators and natural enemies that serve to support crop production (5, 6). There has been a concomitant effort to reduce reliance on insecticides and/or minimize their non-target effects, including state and federal regulation, Integrated Pest Management (IPM) programs, and the development of alternative pest-management technologies and production/marketing systems such as organic farming (7).

Nonetheless, studies suggest recent and widespread declines in insect abundance, diversity, and range (8-10), and insecticide use has been identified as a likely contributor along with habitat loss, species introductions, and climate change (11). In the US, declines have been documented in populations of several wild bee species, butterflies in Ohio and lowland California, and the migratory monarch butterfly (12-15), while beekeepers sustain losses of > 40% of their managed honey bee colonies annually (16). There is also evidence that populations of some insect pests are declining (17, 18).

To understand trends in pest management and potential impacts on target and non-target insect species, we argue it is important to disentangle four distinct dimensions of agricultural insecticide use, organized into two broad categories (Figure 1), extent and intensity. At the landscape scale, the *extent* of insecticide use is a function of the area devoted to cropland as well as the proportion of that cropland treated with insecticides. Historically in the US, a majority of cropland has not been treated with insecticides (19), likely because insect pest populations have not been problematic enough to justify treatment in low-value, large-acreage commodity crops such as wheat, soybean, and corn. For those fields that are treated, the *intensity* of insecticide use varies as a function of the application rate (kg/ha) of products applied and the potency (toxicity/kg) of those products toward insects, typically measured via the LD_50_ (the dose required to kill 50% of a population). These factors jointly determine *insect toxic load*, the total number of insect lethal doses applied in a given area (as described in 20).

**Figure 1.**
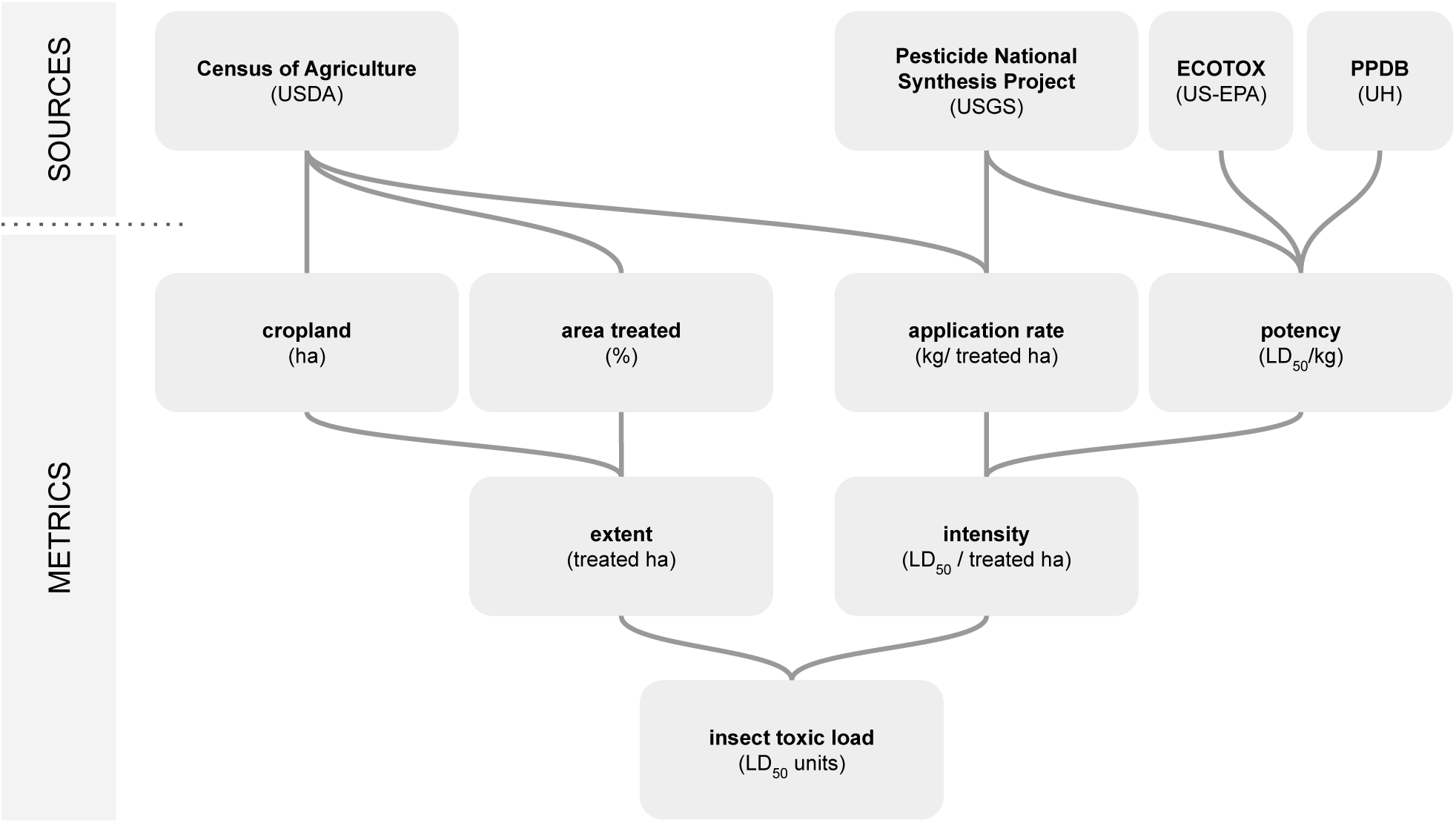
Schematic of insect toxic load, its drivers, and the data sources used to generate these values in this analysis.

Past analyses of insecticide use on US farmland, focusing on only a limited subset of the contributors to insect toxic load, have arrived at divergent narratives of overall trends in insecticide use. For example, analyses based on farmer survey data from the US Department of Agriculture (USDA) show long-term national declines in the application rate of insecticides in corn and cotton crops, and corresponding reductions in the overall weight of insecticides applied (19, 21). A common interpretation of these trends is that introduction of transgenic insect-resistant (*Bt*) crops starting in the mid-1990s led to a reduction in insecticide application (19, 21). Conversely, data from the US Agricultural Census indicate that the extent of US insecticide use has *increased* since 1997, precisely in those regions dominated by *Bt* corn production (22-24). The hypothesized cause of this increase in extent is the widespread adoption of neonicotinoid seed treatments (23, 25). The first analysis did not consider insecticide potency, while the second did not account for intensity. As these examples illustrate, the use of different metrics and data sources leads researchers to conflicting interpretations of overall trends in insecticide use and their associated drivers.

Importantly, describing aggregate trends in pesticide use is not very meaningful without a consideration of the potency of diverse active ingredients (20, 26). In the US alone, there are more than 1000 pesticide active ingredients registered for use (27), which vary by orders of magnitude in potency to target and non-target organisms (28). Aggregate trends in application rate or total weight applied can mask important shifts in the use of different insecticide products over time. For insecticides, this is particularly problematic given evidence of a long-term trend toward development of active ingredients with increasing potency toward insects (29).

Finally, insecticide use trends can vary substantially depending on the spatial scale considered. Analyses at the national scale may obscure important local variation, particularly in the US given the large size of the country, the concentration of different crop production systems in particular areas, the divergent insecticide regimes associated with different crops, and regional variation in pest pressure (24, 30, 31). Analyses at regional to local scales can help to target insecticide mitigation efforts and facilitate research relating spatial patterns in insecticide use to spatial patterns in important outcomes such as crop production, insecticide resistance, and pollinator decline. Given limitations of existing data sources, most previous studies at the county scale have focused solely on the extent of insecticide use, missing potentially important changes in intensity.

Here, we integrate data from several national databases on the components of insecticide extent and intensity (Figure 1) to characterize spatiotemporal patterns in insect toxic load and its drivers from 1997-2012 at the county scale. We use the honey bee (*Apis mellifera*) as a representative insect species to calculate insect toxic load because it is the standard terrestrial insect in regulatory pesticide tests, and so has the most comprehensive toxicity data available. Analyses based on both contact toxicity (exposure to the outside of the body) and oral toxicity (exposure via ingestion) captures uncertainty and variability in exposure pathways. Our goals were to *i*) assess trends in insect toxic load on a contact and oral basis from 1997 to 2012, nationally and for counties in the contiguous US, *ii*) identify the relative contribution of factors (extent and intensity) responsible for these changes over the same time period, and *iii*) describe regional variation in these trends among agricultural production regions.

## Results

### Trends in insect toxic load

Between 1997 and 2012 insect toxic load across the contiguous US increased more than 9-fold on an oral LD_50_ basis and was roughly constant on a contact LD_50_ basis, despite an overall decline in the weight of insecticides applied (Figure 2). Toxic load was spatially heterogenous (Figure 3); in 2012, the top 10% of counties accounted for 55% and 48% of contact and oral toxic load, respectively. From 1997 to 2012, oral toxic load increased in 87% of counties and decreased in 13% of counties (median change: +15-fold, IQR: +3.2-fold to +103-fold) while contact toxic load increased in 51% of counties and decreased in 49% of counties (median change: +5%, IQR: −56% to +142%).

**Figure 2.**
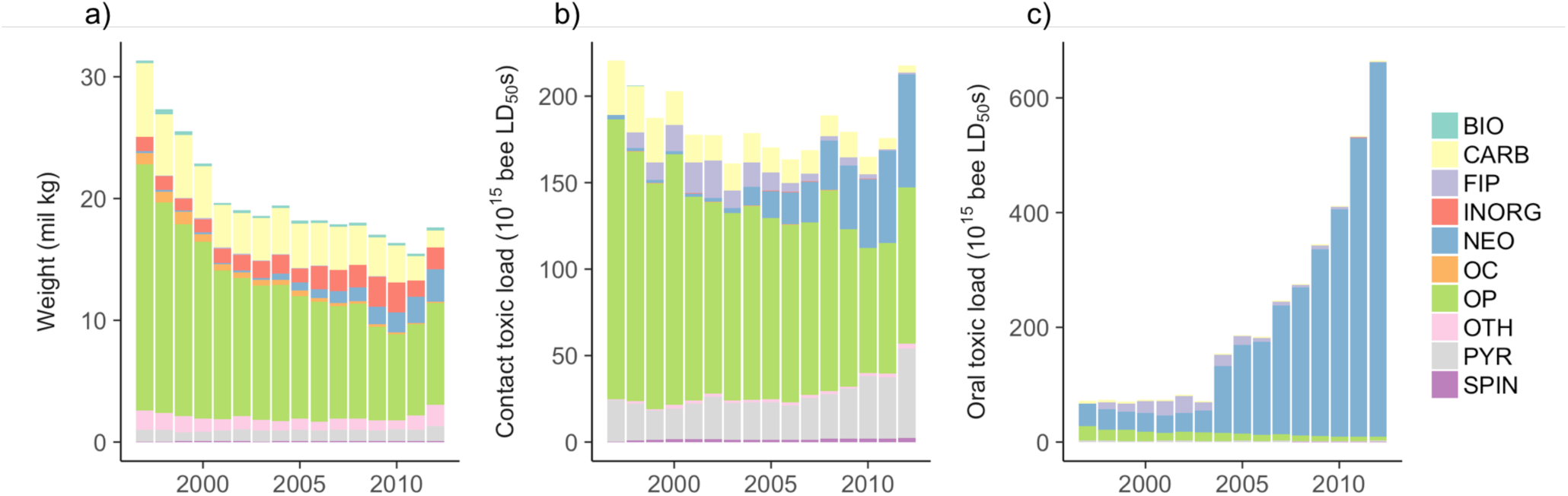
National trends in weight (a), and insect toxic load of all agricultural insecticides applied on a contact (b) and oral (c) toxicity basis from 1997 to 2012 by insecticide class, for the contiguous US. Mann-Kendall tests indicated a monotonic decrease in weight (tau = −0.88, *P* < 0.001), insignificant change in contact toxic load (tau = −0.27, *P* = 0.16), and a monotonic increase in oral toxic load (tau = 0.85, *P* < 0.001). Key: BIO = biological, FIP = fipronil, NEO = neonicotinoid, SPIN = spinosad, PYR = pyrethroid, CARB = carbamate, OP = organophosphate, OC = organochlorine, INORG = inorganic, and OTH = other.

**Figure 3.**
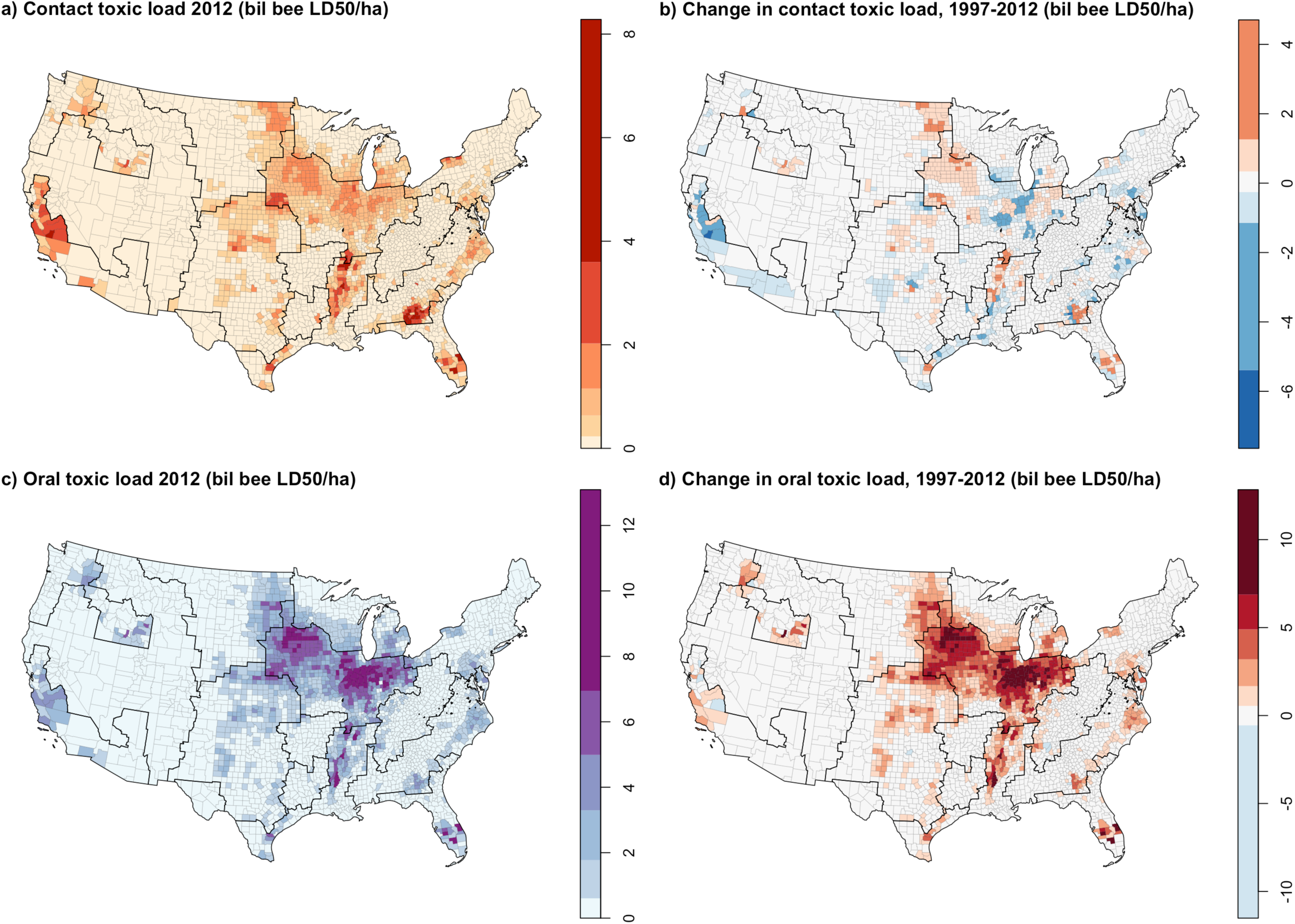
County-level patterns in contact (a, b) and oral (c, d) insect toxic load of agricultural insecticides in 2012 (a, c) and fold change from 1997-2012 (b, d). Toxic load is normalized by the total area (ha) of each county, to account for variable county size. Note that California data excludes seed-applied insecticides. Color scales are based on Jenks natural breaks, which maximize similarity within classes and minimize similarity between classes, adjusted for (b) and (d) to center on zero. Black outlines show USDA Farm Resource Regions.

In 2012, we estimate that agricultural insecticides were applied to ∼5% of the land area of the contiguous US (26% of cropland), at an average intensity of 5 billion honey bee contact LD_50_s per treated ha and 16 billion honey bee oral LD_50_s per treated ha (Table 1).

**Table 1.**
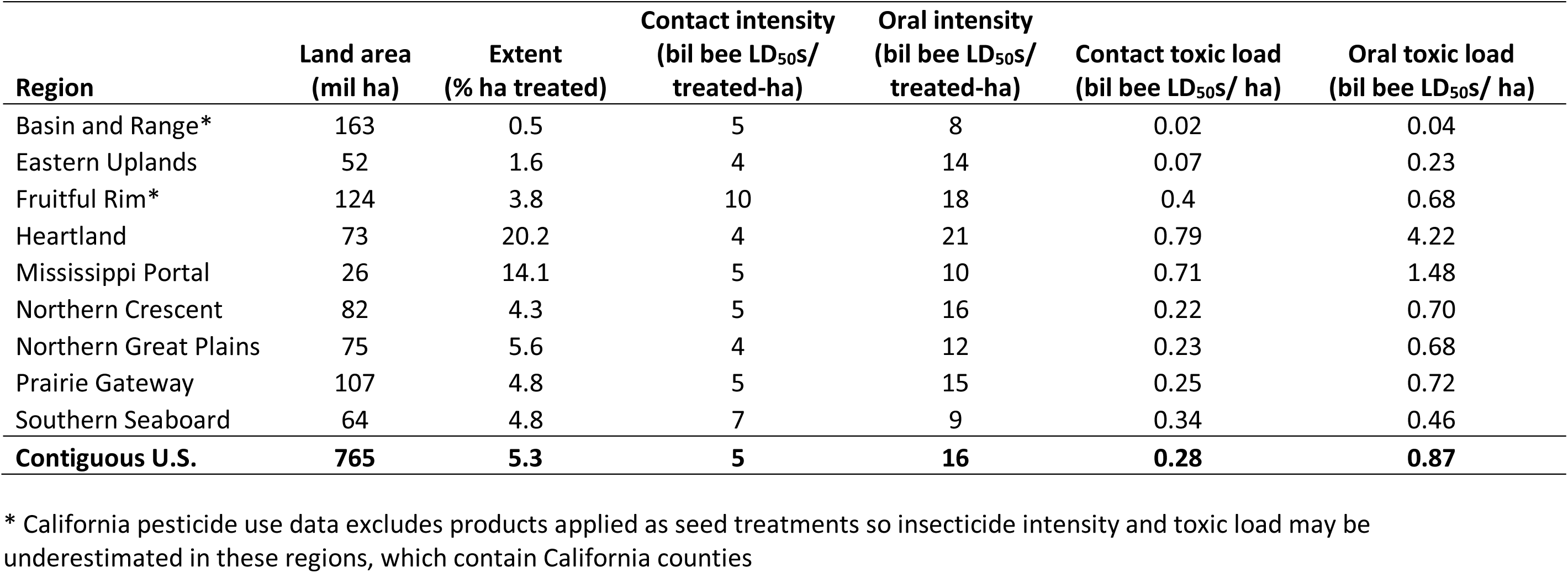
Estimates of 2012 insect toxic load and its contributors associated with insecticide use in the nine USDA Farm Resource Regions of the contiguous US.

| Region | Land area<br>(mil ha) | Extent<br>(% ha treated) | Contact intensity<br>(bil bee LD<sub>50</sub>s/<br>treated-ha) | Oral intensity<br>(bil bee LD<sub>50</sub>s/<br>treated-ha) | Contact toxic load<br>(bil bee LD<sub>50</sub>s/ ha) | Oral toxic load<br>(bil bee LD<sub>50</sub>s/ ha) |
| --- | --- | --- | --- | --- | --- | --- |
| Basin and Range* | 163 | 0.5 | 5 | 8 | 0.02 | 0.04 |
| Eastern Uplands | 52 | 1.6 | 4 | 14 | 0.07 | 0.23 |
| Fruitful Rim* | 124 | 3.8 | 10 | 18 | 0.4 | 0.68 |
| Heartland | 73 | 20.2 | 4 | 21 | 0.79 | 4.22 |
| Mississippi Portal | 26 | 14.1 | 5 | 10 | 0.71 | 1.48 |
| Northern Crescent | 82 | 4.3 | 5 | 16 | 0.22 | 0.70 |
| Northern Great Plains | 75 | 5.6 | 4 | 12 | 0.23 | 0.68 |
| Prairie Gateway | 107 | 4.8 | 5 | 15 | 0.25 | 0.72 |
| Southern Seaboard | 64 | 4.8 | 7 | 9 | 0.34 | 0.46 |
| **Contiguous U.S.** | **765** | **5.3** | **5** | **16** | **0.28** | **0.87** |

\* California pesticide use data excludes products applied as seed treatments so insecticide intensity and toxic load may be underestimated in these regions, which contain California counties

### Drivers of insect toxic load

The main driver of the increase in oral toxic load was insecticide intensity: a 16-fold increase in oral potency far surpassed a 64% decline in application rate (Figure 4). The increase in insecticide potency was associated with the shifting composition of insecticide classes, away from organophosphates and toward pyrethroids and neonicotinoids (Figure 2, Figure S1); by 2012 neonicotinoids alone contributed 98% of oral toxic load. While contact potency also increased from 1997 to 2012 (+76%), this change was roughly counterbalanced by a corresponding decline in application rate.

**Figure 4.**
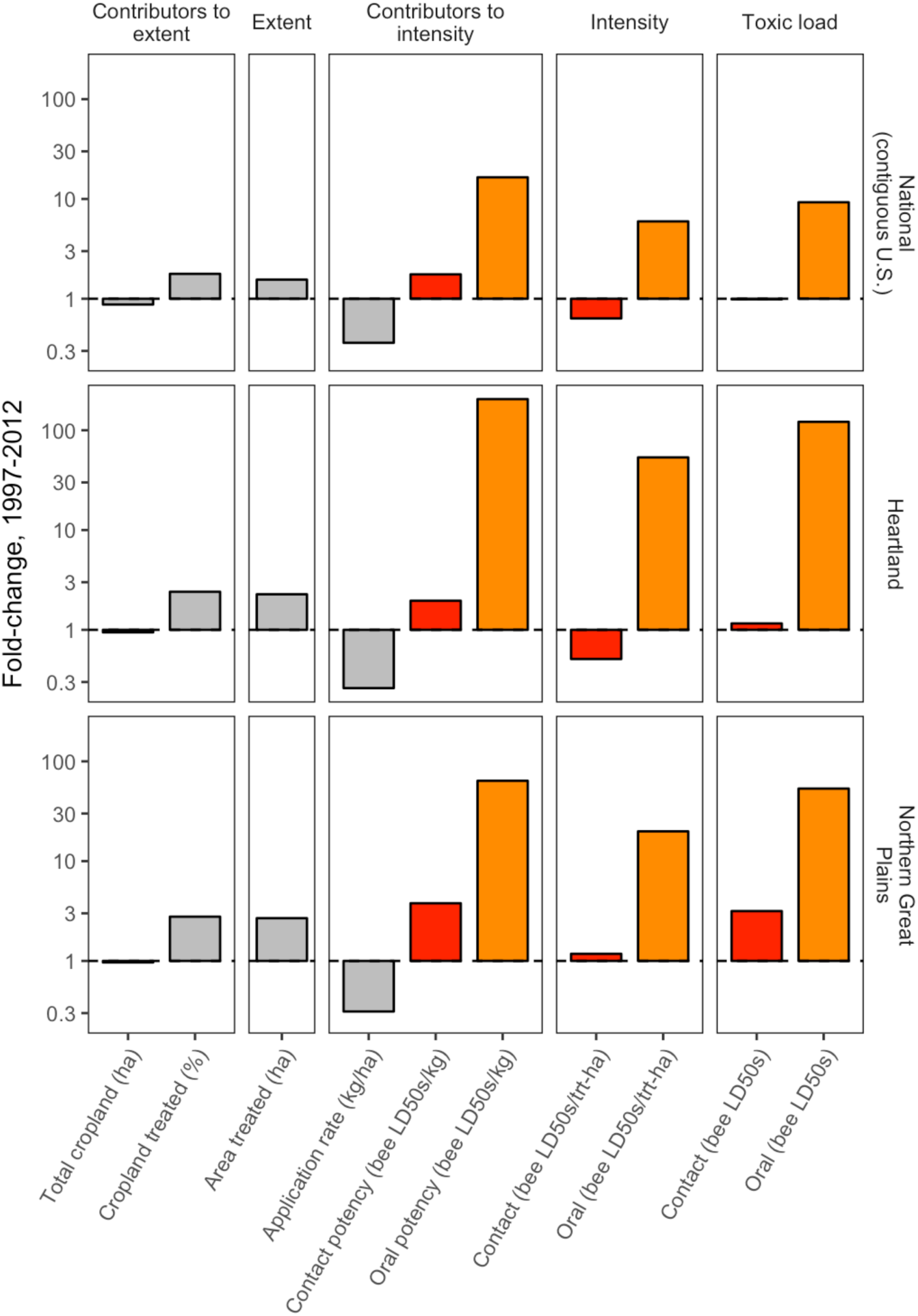
Change in the drivers of insect toxic load from 1997 to 2012 for the contiguous U.S. and select USDA Farm Resource Regions. Fold-change is calculated as a response ratio: Value_2012_/Value_1997_, so that a value of one represents no change, three represents a tripling, 0.3 represents a 70% decline, etc. (values are presented on a log scale). Red represents calculations from contact toxicity and orange from oral toxicity.

Changes in extent also occurred over the study period and influenced overall toxic load; while total cropland declined slightly (−12%), the proportion of cropland treated with insecticides increased by 78%, from 15% of cropland treated in 1997 to 26% in 2012.

### Regional variation in insect toxic load and its drivers

The temporal dynamics of insect toxic load varied across the 9 major agricultural production regions (Figures 3-5). Oral toxic load increased significantly in all regions except the Basin and Range (Table S1), with reductions in application rate overwhelmed by increases in potency and cropland treated (Figure S3). The most dramatic increases occurred in the Heartland (121-fold increase) and the Northern Great Plains (10-fold increase) (Figures 4-5). This pattern was underscored by time series cluster analysis, where the majority of total variation was captured in the node dividing the Heartland + Northern Great Plains cluster from the other seven regions (Figure 5). The cause of this pattern, evident when oral toxic load is parsed by chemical class, was the substantial application of neonicotinoid insecticides in the Heartland and Northern Great Plains, beginning in 2006 and increasing exponentially through 2012 (Figure 5).

**Figure 5.**
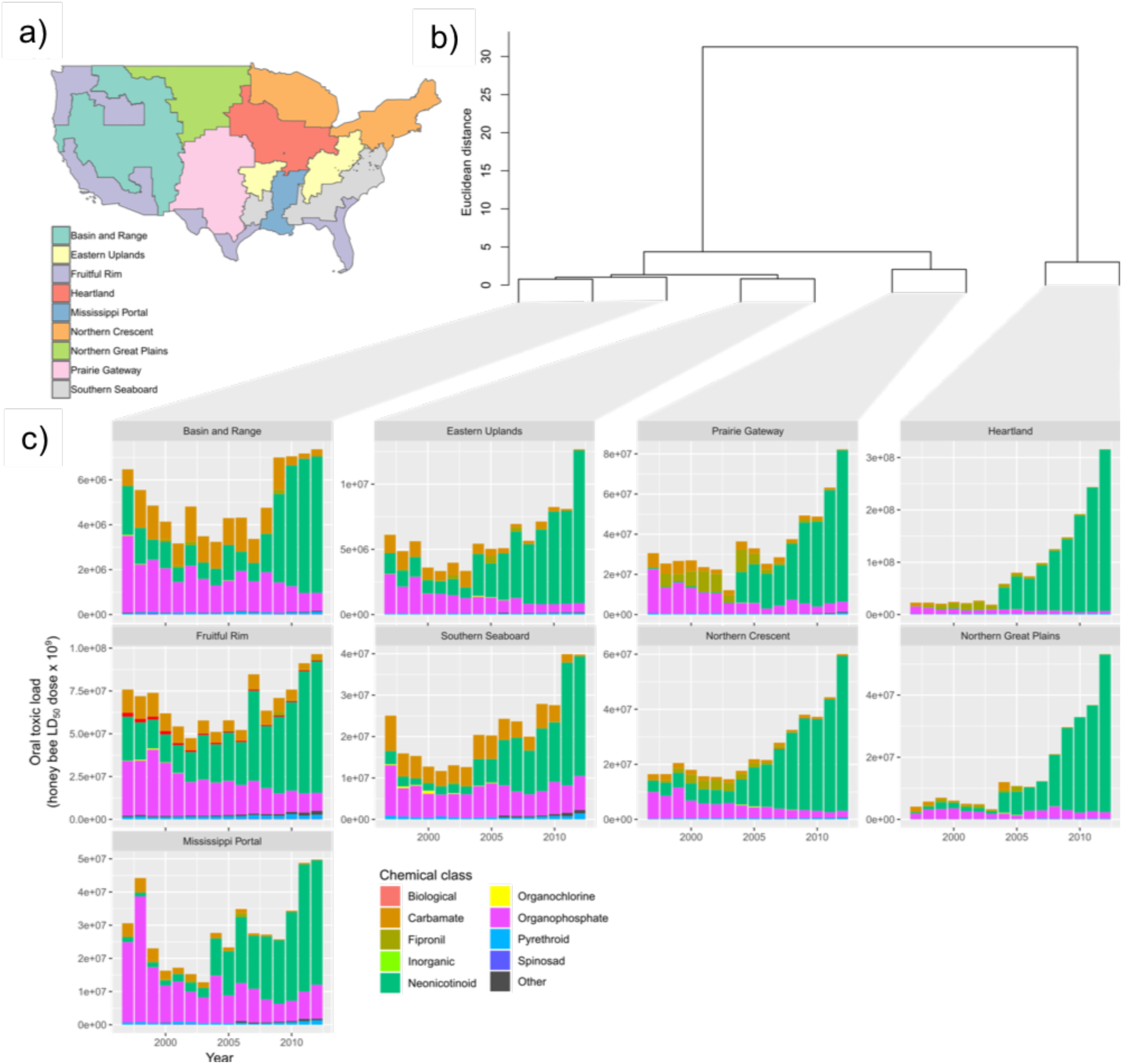
Oral toxic load by region and chemical class. Toxic load time series were constructed for each of the nine USDA Farm Resource Regions (a). Hierarchical clustering (b) grouped regions with similar patterns of toxic load using a Euclidean distance matrix and Ward’s linkage method. The y-axis distance between two nodes and their nearest common node is inversely proportional to similarity. Hence, the majority of variation among regions is captured in the first split that separates the Heartland and Northern Great Plains from the other seven regions. The contribution of different chemical classes to the overall toxic load pattern in each region is depicted with stacked bar plots (c).

Conversely, changes in contact toxic load were more modest and variable, with most regions experiencing no significant trends (Table S1). The exceptions included the Eastern Uplands and Northern Crescent, where significant declines were observed (−38% and −10%, respectively), driven by application rate, and to a lesser extent, declines in total cropland (Table S1, Figure S3). Conversely, contact toxic load increased more than 3-fold in the Northern Great Plains (Table S1, Figure S3), a pattern driven by a 2.7-fold increase in cropland treated and a 3.8-fold increase in contact potency (Figure 4). In cluster analysis, the majority of variation was captured in the node dividing the Northern Great Plains region from all other regions (Figure S4). Notably, organophosphate toxic load increased in the Northern Great Plains from 2006 to 2012, in contrast to all other regions where the contribution of this chemical class was stable or declining (Figure S4).

### Sensitivity analysis

We performed several analyses to characterize the robustness of our results to uncertainty in the underlying data sources and interpolation of missing values (Supplemental Material). We found that interpolation of insecticide use, crop area, and treated area in counties with missing values in our source data had only a minor effect on the overall results (Figures S5-S7). Furthermore, while the LD_50_ values for some of the insecticides used in our analysis were derived from sources with low and medium certainty, the vast majority of insect toxic load was derived from insecticides with LD_50_ values reported in high quality sources (Figure S8). Finally, while we used the more conservative USGS ‘low’ pesticide estimate in our analyses, repeating the analyses with the ‘high’ estimate did not influence our conclusions regarding national and regional trends, although numerical estimates for contact and oral toxic load differed (Figure S9, Table S2).

## Discussion

By creating a novel dataset synthesizing land use, insecticide use, and honey bee toxicity, we characterized spatiotemporal patterns in the total inputs of insecticides to cropland in the contiguous US. We found that insect toxic load from 1997 to 2012 was relatively constant when calculated using contact LD_50_ values, but increased nine-fold when calculated using oral LD_50_ values, consistent with a recent analysis from the United Kingdom (20). Insect toxic load increased despite a decrease in average application rate, which emphasizes the importance of accounting for potency when assessing trends in insecticide use, as recently argued for herbicide use (26, 32). The pattern of divergence between decreasing application rate and increasing extent and potency, combined with significant variation across agricultural regions, helps to explain and reconcile the conflicting narratives of insecticide dependency that characterize recent research on US insecticide use trends (19, 21-24, 32).

Insect toxic load – calculated on an oral LD_50_ basis – increased in all regions, but the increase was particularly acute in the Heartland and Northern Great Plains regions. We attribute this pattern to the increasing use of neonicotinoid seed treatments in corn and soybean. In the Heartland, > 90% of cropland (excluding pasture) in 1997 and 2012 was devoted to these two crops (Figure S2), while the Northern Great Plains experienced significant land use change over the study period in which corn and soybean displaced other crops and conservation land (33). Neonicotinoids accounted for the overwhelming majority of oral toxic load by 2012, and previous research (25) showed that virtually all neonicotinoid use in corn and soybean is via seed application.

The important role of neonicotinoid seed treatments in this analysis underscores another source of conflicting interpretations of US insecticide use trends – inconsistent or incomplete data collection. USDA started collecting data on seed treatments only in 2015 (34), and the data are not yet publicly available. The USGS Pesticide National Synthesis Project included seed-applied pesticides through 2014, then stopped when the data provider excluded them (35). Analyses derived from the Census of Agriculture suggest that seed treatments are at least partly captured by that dataset (23, 24), although the survey language is ambiguous (36). Consequently, our estimate of insecticide extent is likely an underestimate. Our results show that analyses of insecticide use that do not account for seed treatments are likely missing a major component of insect toxic load, highlighting a need for more consistent and comprehensive data collection. Similarly, our results point to a need for more detailed investigation of the combined agronomic and socioeconomic drivers of neonicotinoid seed treatments, which do not seem to be cleanly related to pest pressure or field-level economics (25, 31).

Recommendations to farmers for reducing exposure of pollinators and natural enemies to insecticides have historically emphasized avoiding contact exposure, for example by spraying at night or not spraying during bloom (e.g. 6). Our finding that oral toxic load - driven by the use of systemic neonicotinoids - has surpassed contact toxic load suggests that this guidance may need to be amended. Systemic insecticides are taken up by plant tissues and can remain active for long periods. Indeed, a major route of exposure for foraging bees to neonicotinoids and other pesticides through nectar and pollen obtained from flowering weeds in agricultural areas, where the weeds have taken up residues from the soil (37-39). Seed treatments likely exacerbate this route of exposure, since neonicotinoids can remain present in the soil for years; indeed, recent studies identified imidacloprid residues in plants and bee colonies, though imidacloprid use had been discontinued in these study regions in previous years (37, 40). It is sometimes possible to adjust management practices to avoid non-target exposure to systemic pesticides, but this requires longer-term planning than mitigation at the time of spraying. In apples, for example, spraying neonicotinoids 5-10 days prior to bloom ensures levels are sufficiently low to protect pollinators and natural enemies (41).

Importantly, our approach is not a formal risk assessment, nor does it fit neatly into existing risk assessment frameworks (e.g. 42). The formal concept of risk--the likelihood of adverse outcome--is defined as the product of hazard and exposure. Insect toxic load corresponds to the hazard component of this definition, but we do not attempt to estimate or model pesticide fate in the environment or exposure. Formal risk assessments are also, by necessity, narrow in scope, typically evaluating single compounds with respect to specific receptor organisms over a narrow range of use scenarios. In contrast, our approach integrates across compounds and use scenarios to estimate cumulative, landscape-scale insecticide hazard to insects in general, represented by a honey bee proxy. In this sense, we present an insect “hazard cup”, analogous to the “risk cup” paradigm used in human risk assessment (43) and echoing Berenbaum’s (44) extension of this paradigm to honey bee risk assessment.

The development of a county-level measure of insect toxic load, aggregated across insecticides and national in scope, advances our ability to evaluate the impacts of insecticide use on target and non-target insect populations. Previous large-scale studies have relied on the percentage of land in agriculture (45), the percentage of land used to produce particular crops (46), the percentage of land treated (23, 47), the weight of insecticides applied (48), or weight of specific compounds (14, 49). By developing a pipeline that integrates information from national databases to summarize insecticide intensity, extent, and insect hazard (toxic load), our study provides a common and repeatable framework that can be used across the US to investigate the potential impacts of insecticide use on diverse insect populations and dependent wildlife. Furthermore, these national maps of insect toxic load, in combination with maps examining insect abundance and ecosystem services (50), can be used to identify regions that should be prioritized for conservation (areas where insect abundance, diversity, or ecosystem services are high and insecticide hazard is low) or mitigation (areas where abundance of imperiled species, such as monarch butterflies, is high (51) and insecticide hazard is also high). While there are clearly limitations to this analysis, it complements more traditional ‘bottom-up’ approaches to ecotoxicology by enabling ‘top-down’ research at the landscape scales at which insect populations and ecosystem services are structured (52).

## Materials & Methods

This study required synthesizing data on insecticide use, land use, and toxicity from multiple national databases to generate a novel dataset documenting county-level spatiotemporal patterns in insect toxic load and its drivers. All data processing and analysis was performed in the R statistical language, version 3.6.1 (53). The code used to generate and analyze the dataset is stored on GitHub and the workflow will be made publicly available upon publication.

### Insecticide use data

Data on the weight of insecticides applied by county and year were downloaded from the US Geological Survey (USGS) Pesticide National Synthesis Project (54). This dataset reports pesticide use (kg active ingredient applied in each county) for > 400 of the most commonly applied herbicides, insecticides, and fungicides on agricultural crops. The estimates are derived from proprietary farmer surveys, except for California, where estimates are drawn from the state’s Department of Pesticide Regulation (54). The dataset includes seed-applied pesticides through 2014, except in California, where they have never been included. USGS provides two pesticide estimates; we used the more conservative ‘low’ estimate (data presented in the text and figures) but repeated our analyses with the ‘high’ estimate (data presented in the Supplementary Materials) to ensure they were robust to differences in pesticide estimation. We focused on the 138 insecticides in the dataset, classified as such based on the Pesticide Properties Database (PPDB; 55) and the Insecticide Resistance Action Committee (56). Following the custom of the US Environmental Protection Agency (US-EPA) we excluded petroleum oils and distillates (4).

### Land use data

The US Census of Agriculture is conducted every five years. We used this dataset as a source of county-level data for the years 1997, 2002, 2007, and 2012 on *i*) cropland area treated with at least one insecticide, *ii*) total cropland area, and *iii*) total land area in each county. Data were downloaded from the developer page of the Quick Stats database maintained by the National Agricultural Statistics Service (57). We focused on area in cropland rather than pastureland because USGS data indicates that only ∼1% of insecticides are applied to pastureland. Missing values occurred in the dataset when data were withheld to avoid disclosing information about individual farm operations; this situation occurs mostly in cases where agricultural acreage in a county is very low. In those cases, we imputed missing values with the average of values in other years for that county if possible, or with zeros if values for the county were missing in all years (24). Finally, counties were grouped into regions using a system developed by US Department of Agriculture (‘Farm Resource Regions’) to capture geographic variation in farming systems related to cropping patterns, similarities in soil and climate, and economic characteristics (30).

### Toxicity data

We considered toxicity on the basis of contact and oral exposure. Contact exposure occurs when an insect encounters the insecticide on the outside of its body, for example if it is directly sprayed. Oral exposure occurs when an insect ingests the insecticide, for example when feeding on a recently treated plant. We conducted our analyses separately with each toxicity measure because of the difficulty of predicting the dominant mode of exposure from insecticide use data alone (particularly given that the data do not include method of application).

Honey bee acute toxicity values were compiled mainly from US-EPA’s ECOTOX database (58) and the PPDB. The ECOTOX database was queried in July 2017 and values were recorded from the PPDB in June 2018. Both sources include toxicity data generated during pesticide regulatory procedures as well as from other sources (e.g. scientific journal articles). We gave preference to values generated in the regulatory processes of the US-EPA and the European Food Safety Authority (EFSA) because they tend to be generated using standardized testing protocols.

To compile honey bee LD_50_ values for the 138 insecticides present in the USGS dataset, we first searched ECOTOX for all lab-generated LD_50_ estimates for *Apis mellifera*. We then processed the data by recoding exposure types into ‘contact’, ‘oral’, or ‘other’ and standardizing units where possible into ug/bee, consulting original sources as needed to verify extreme values and to fill in missing information. The following criteria were used to select records: *i*) exposure time 4 days or less (e.g. acute, after 59), *ii*) contact or oral exposure, *iii*) tests on adult bees, and *iv)* non-zero LD_50_ estimate in units of µg/bee. We related this cleaned dataset to USGS pesticide names on the basis of CAS numbers. Similarly, we generated a dataset comprising honey bee LD_50_ values (acute contact and oral) from the PPDB, along with the source code identifying high-quality values coming from the EU regulatory process (code ‘A5’).

To generate a consensus list of contact and oral LD_50_ values for all insecticides reported in the USGS dataset (Supplemental Data 1) we used the following procedure (adapted from 60):

1. If point estimate(s) were available from US or EU regulatory bodies, we used those values, taking a geometric mean when estimates were available from both sources; otherwise,
2. If point estimate(s) were available from other sources in ECOTOX or PPDB, we used those values, taking a geometric mean when estimates were available from multiple sources; otherwise,
3. If unbounded estimate(s) were available from US or EU regulatory bodies (i.e. “greater than” or “less than” some value), we used the minimum (for “less than”) or the maximum (for “greater than”); otherwise,
4. If unbounded estimate(s) were available from other sources in ECOTOX or PPDB, we used the minimum (for “less than”) or the maximum (for “greater than”); otherwise,
5. We used the median toxicity value for the insecticide mode-of-action group, with group defined by the Insecticide Resistance Action Committee.
6. In rare cases (n = 1/138 compounds for contact toxicity and 8/138 compounds for oral toxicity) after this procedure we were still left without a toxicity estimate for a particular compound. In those cases, we used the median for all insecticides.

### Data synthesis and analysis

We synthesized the data sources described above to generate a novel dataset describing insect toxic load and its contributors (Supplemental Data 2). First, we calculated the total contact and oral insect toxic load for each county-year combination from 1997 to 2012, using the following equation (after 26):

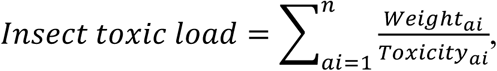

where weight is the total weight (kg) of insecticide active ingredient (*ai*) applied in a county in a particular year, and toxicity is the contact or oral acute toxicity to honey bees (LD_50_ in µg/bee) for each *ai*. There were a minority of counties for which insecticide use data were missing in particular years. We used linear interpolation to fill in missing values for kg applied, contact toxic load, and oral toxic load. If agricultural insecticide use was never reported in a particular county (this occurred in a few, highly urban counties like New York City), we assumed it was zero.

Insect toxic load in a county is a function of the area of land treated with insecticides (*extent*) and the aggregate toxicity of the insecticides applied per unit area (*intensity*) (Figure 1). To better elucidate drivers of insect toxic load over the study period, we calculated elements of insecticide extent and intensity for those years in which land use data were available from the agricultural census (1997, 2002, 2007, 2012). Data on county-level land use were joined with data on insecticide use and toxic load on the basis of county FIPS codes (Supplemental Data 3). Once joined, we identified and calculated the contributors to insect toxic load as defined in Figure 1.

To test for monotonic trends in insect toxic load at the national and regional scales we used the non-parametric Mann-Kendall test (after 26). To further analyze change over time, and to compare the relative magnitude of change across indicators with disparate units, for each of our insecticide indicators we calculated fold-change from 1997 to 2012 as the response ratio: Value_2012_/Value_1997_, so that a value of one represents no change, two represents a doubling, and one half represents a decline of 50%.

To analyze regional differences in the temporal dynamics of toxic load, we performed hierarchical time series clustering using the R package ‘dtwclust’ (61). Contact and oral toxic load were calculated for years 1997-2012 for each USDA Farm Resource Region. Because regions differed by as much as two orders of magnitude in absolute toxic load, we converted the absolute toxic load within each region to fold-change relative to 1997, as described above, enabling more informative clustering driven by patterns of relative toxic load. We then performed hierarchical time series clustering using a Euclidean distance matrix and Ward’s linkage method, with oral and contact toxic load analyzed separately.

## Acknowledgements

This work was supported by the National Socio-Environmental Synthesis Center (SESYNC) under funding received from the National Science Foundation (DBI-1639145) and by grants from USDA-NIFA-AFRI (#2018-67013-27538) and the Foundation for Food and Agricultural Research (#549032). We thank Karan Shakya and Sara Soba for assistance with data processing for LD_50_ values.

## SUPPLEMENTAL MATERIAL

### Sensitivity analysis

We performed several analyses to characterize the robustness of our results to uncertainty in the underlying data sources. One source of uncertainty in our estimates stemmed from interpolating missing values for insecticide use, crop area, and treated area in counties with missing values in our source data. However, the sensitivity analysis revealed that interpolation had a minor effect on estimates of insect toxic load (Figures S5-S7). At the national scale, pesticide use was interpolated for fewer than 2% of counties in each year, and these counties contributed less than 0.05% to weight applied, contact toxic load, and oral toxic load. Interpolation was more frequent for treated cropland; nonetheless cropland treated was interpolated for < 6% of county-year combinations and these interpolated values contributed < 1% to total cropland treated. Across data items, interpolation tended to be rare in the regions driving our results (Heartland, Northern Great Plains), and more common in regions with generally little cropland (e.g. Basin and Range; Figures S5-S7).

Another source of uncertainty in our analyses concerned the honey bee LD_50_ values underlying the translation of insecticide use into insect toxic load. For each US county, we calculated the percentage of insect toxic load contributed by compounds with ‘high’, ‘medium’, and ‘low’ uncertainty in their LD_50_ values. Uncertainty was considered low if LD_50_ values were derived from US or EU regulatory procedures, medium if LD_50_ values were compound-specific but derived from other scientific sources, and high if LD_50_ values were estimated by a median value for the class or insecticides as a whole. In each year of the dataset active ingredients with high-quality values (those generated as part of the US or EU regulatory processes) accounted for the vast majority of insecticide weight (79-86%), contact toxic load (91-98%), and oral toxic load (98-100%; Figure S7). Low-quality values (those missing and so estimated from class medians) contributed modestly to insecticide weight (1.5-15%) but very little to contact toxic load (< 1%) and oral toxic load (< 3%), likely reflecting that active ingredients without honey bee toxicity data are more likely to be found in classes with generally low toxicity.

To test the sensitivity of our findings to different methods of insecticide estimation, we repeated our main analyses using the less-conservative USGS ‘high’ estimates of insecticide use (Figures S3, S8, S9; Table S2). Qualitative patterns in insect toxic load and its drivers were extremely similar for the two estimates, supporting our conclusions regarding overall trends and regional variability in insecticide indicators. However, there were quantitative differences. For example, the 2012 estimate of insect toxic load at the national scale based on the ‘high’ USGS estimate was 39% higher on a contact-toxicity basis and 7% higher on an oral-toxicity basis than the equivalent value using the ‘low’ estimate (Table S2).

## SUPPLEMENTAL FIGURES

**Figure S1.**
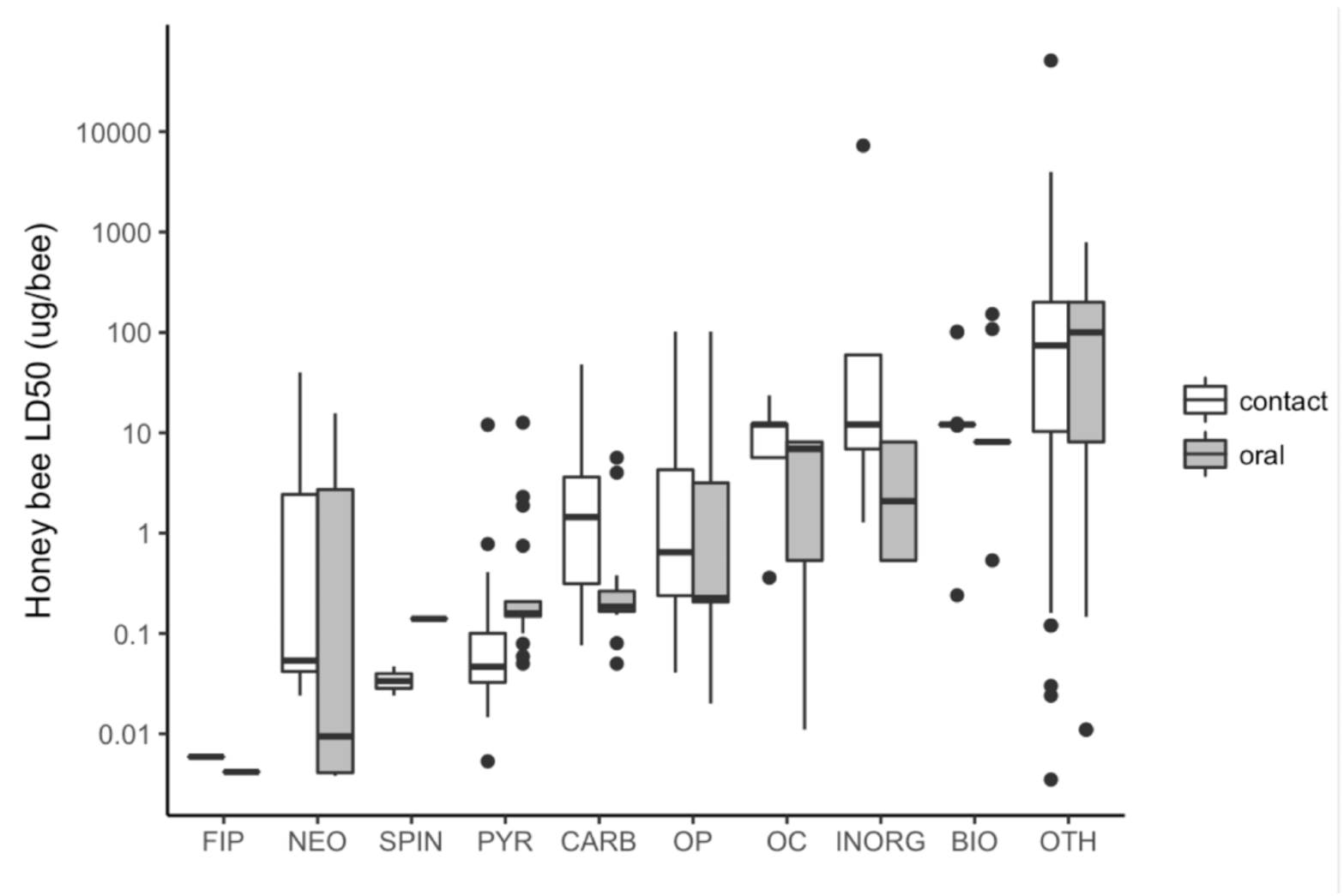
Honey bee LD_50_ values by insecticide class. Key: FIP = fipronil, NEO = neonicotinoid, SPIN = spinosad, PYR = pyrethroid, CARB = carbamate, OP = organophosphate, OC = organochlorine, INORG = inorganic, BIO = biological, and OTH = other.

**Figure S2.**
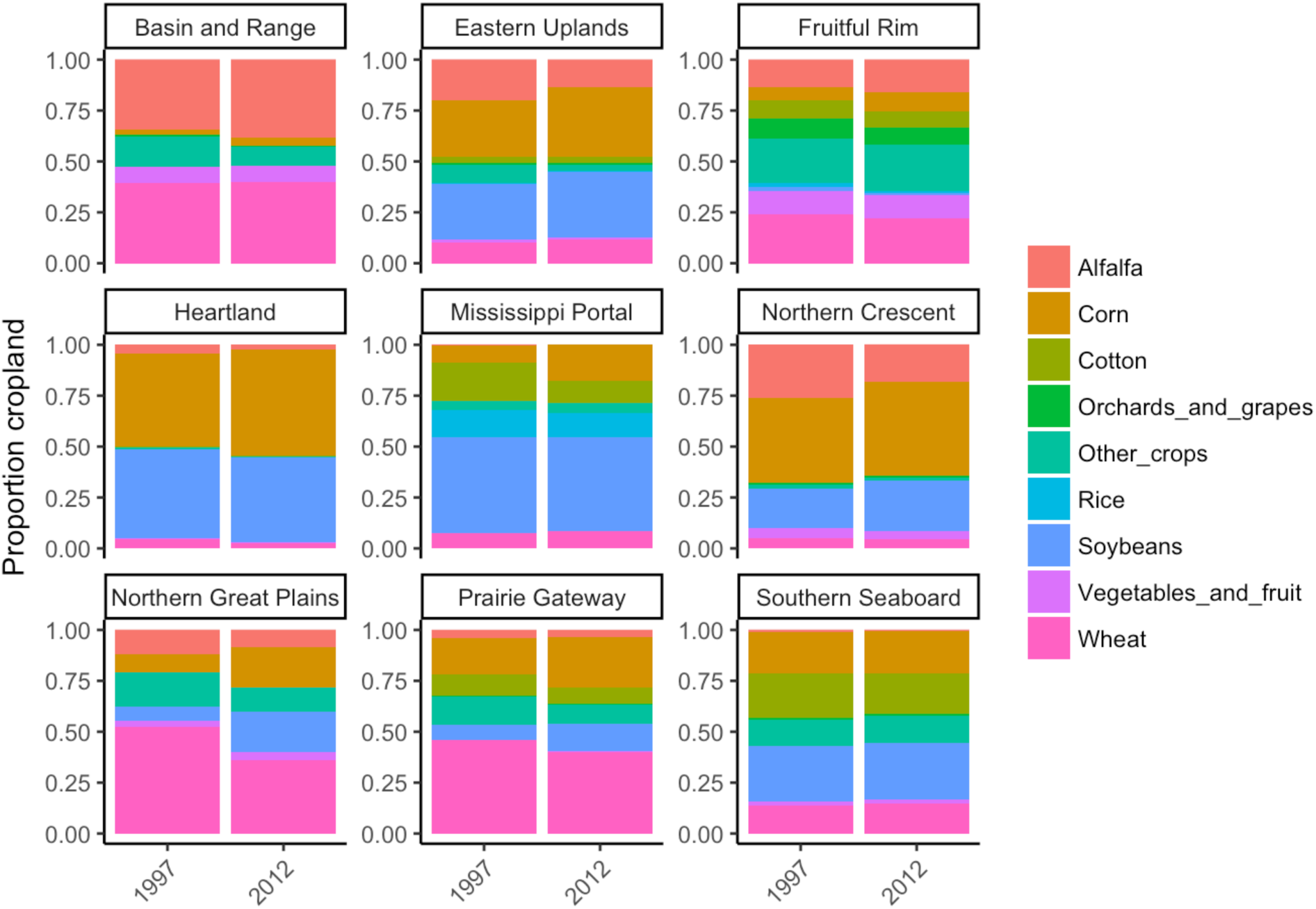
Composition of cropland by region in 1997 and 2012, based on crop area data from the U.S. Census of Agriculture.

**Figure S3.**
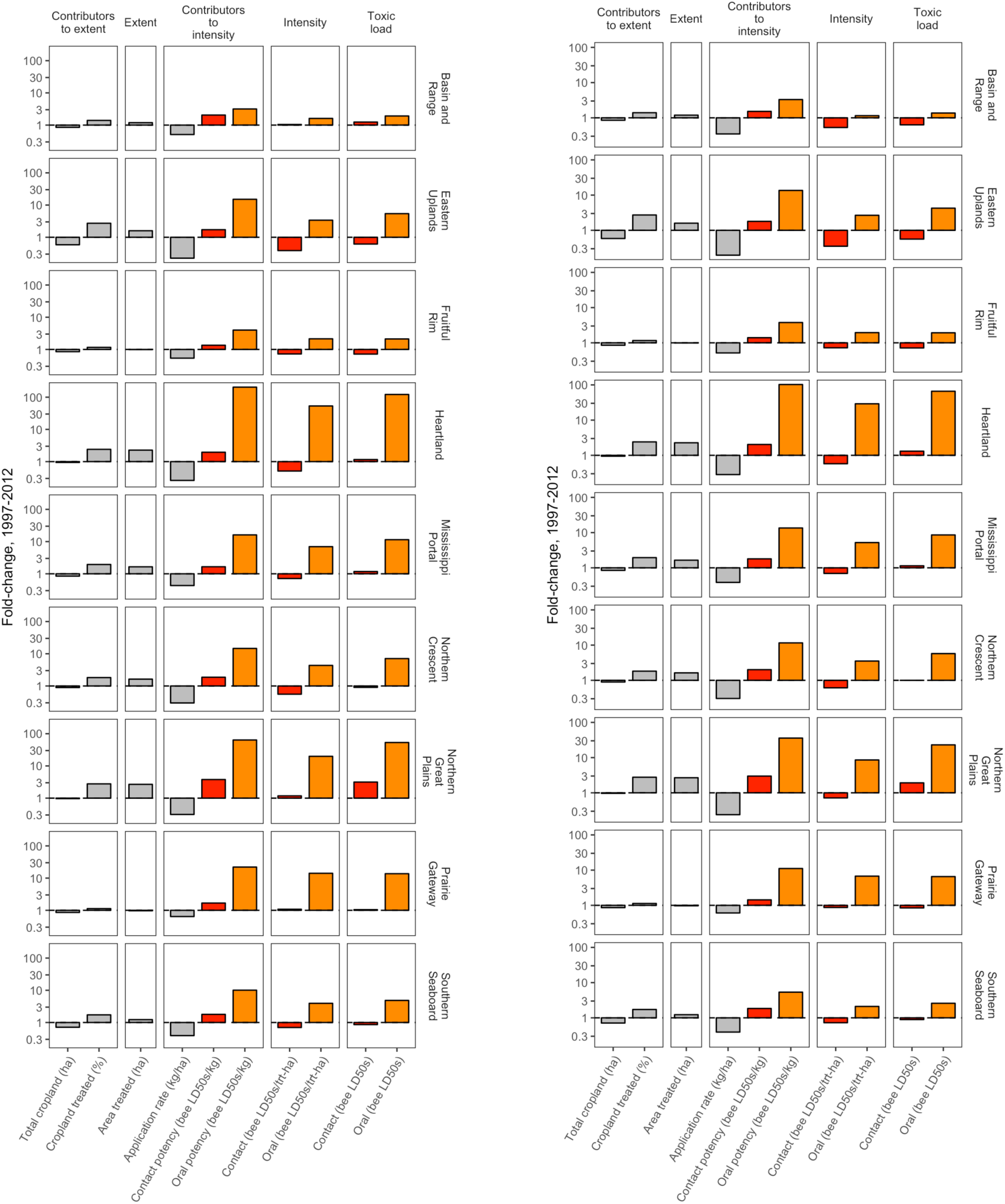
Change in the drivers of insect toxic load from 1997 to 2012 for all agricultural production regions, based on the USGS low (left) and high (right) pesticide use estimates. Fold-change is calculated as a response ratio: Value_2012_/Value_1997_, so that a value of one represents no change, two represents a doubling, one half represents a decline of 50% (values are presented on a log scale).

**Figure S4.**
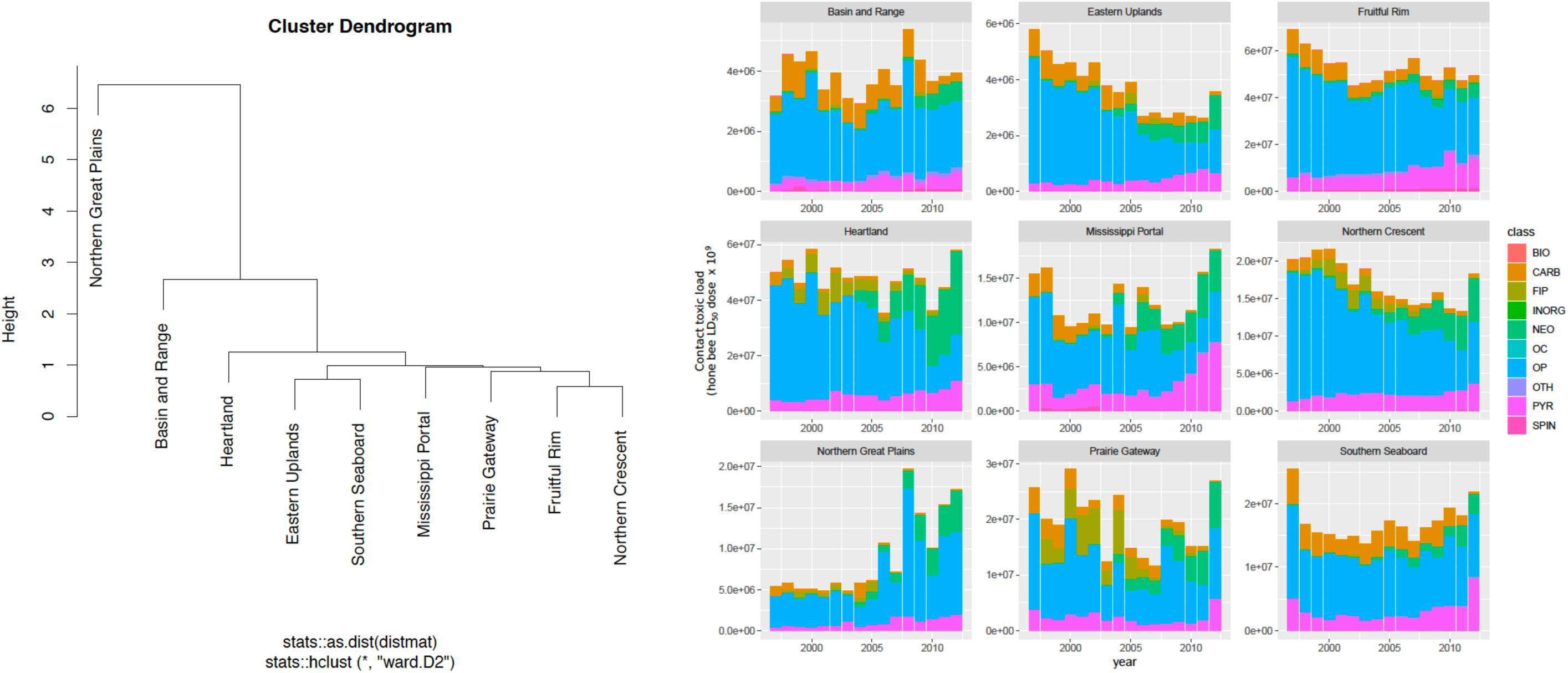
Contact toxic load by region and chemical class. Toxic load time series were constructed for each of the nine USDA Farm Resource Regions. Hierarchical clustering (left) grouped regions with similar patterns of toxic load using a Euclidean distance matrix and Ward’s linkage method. The y-axis distance between two nodes and their nearest common node is inversely proportional to similarity. Hence, the majority of variation among regions is captured in the first split that separates the Northern Great Plains from the other eight regions. The contribution of different chemical classes to the overall toxic load pattern in each region is depicted with stacked bar plots (right).

**Figure S5.**
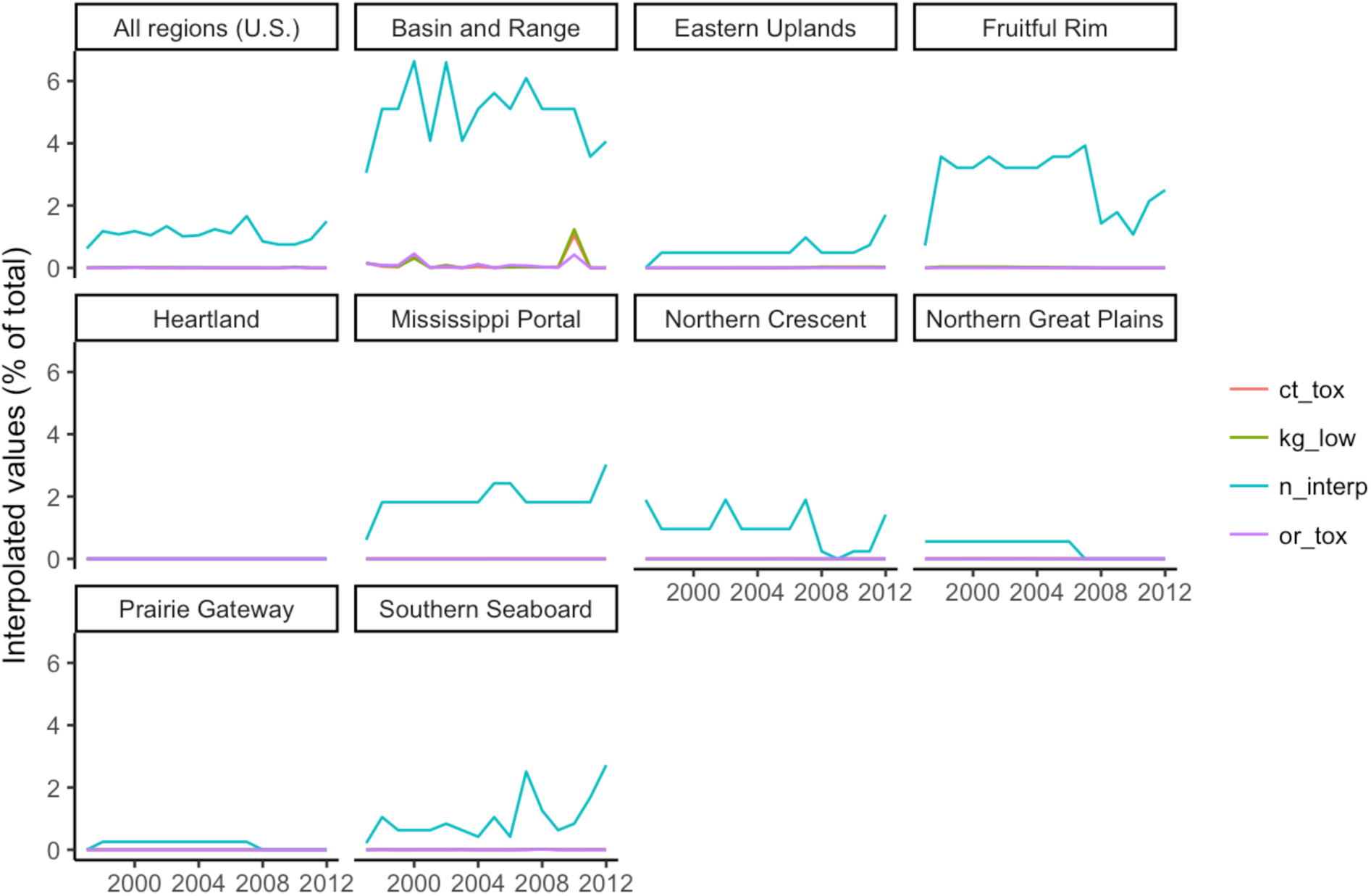
Contribution of counties with interpolated insecticide values to total counties (n_interp), mass applied (kg_low), contact toxic load (ct_tox), and oral toxic load (or_tox) for each of nine agricultural regions and all regions combined.

**Figure S6.**
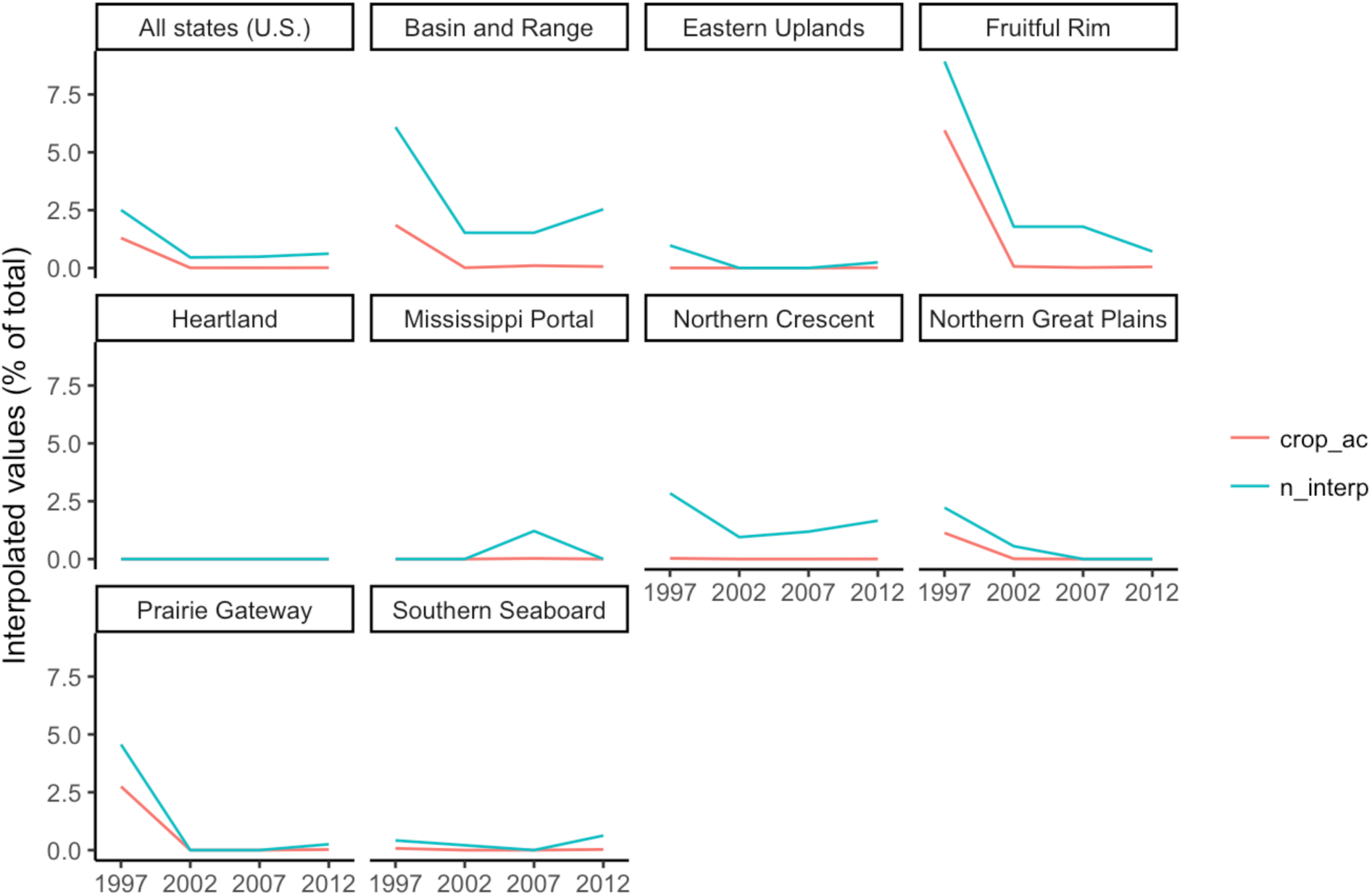
Contribution of counties with interpolated cropland area values to total counties (n_interp) and cropland (crop_ac) for each of nine agricultural regions and all regions combined.

**Figure S7.**
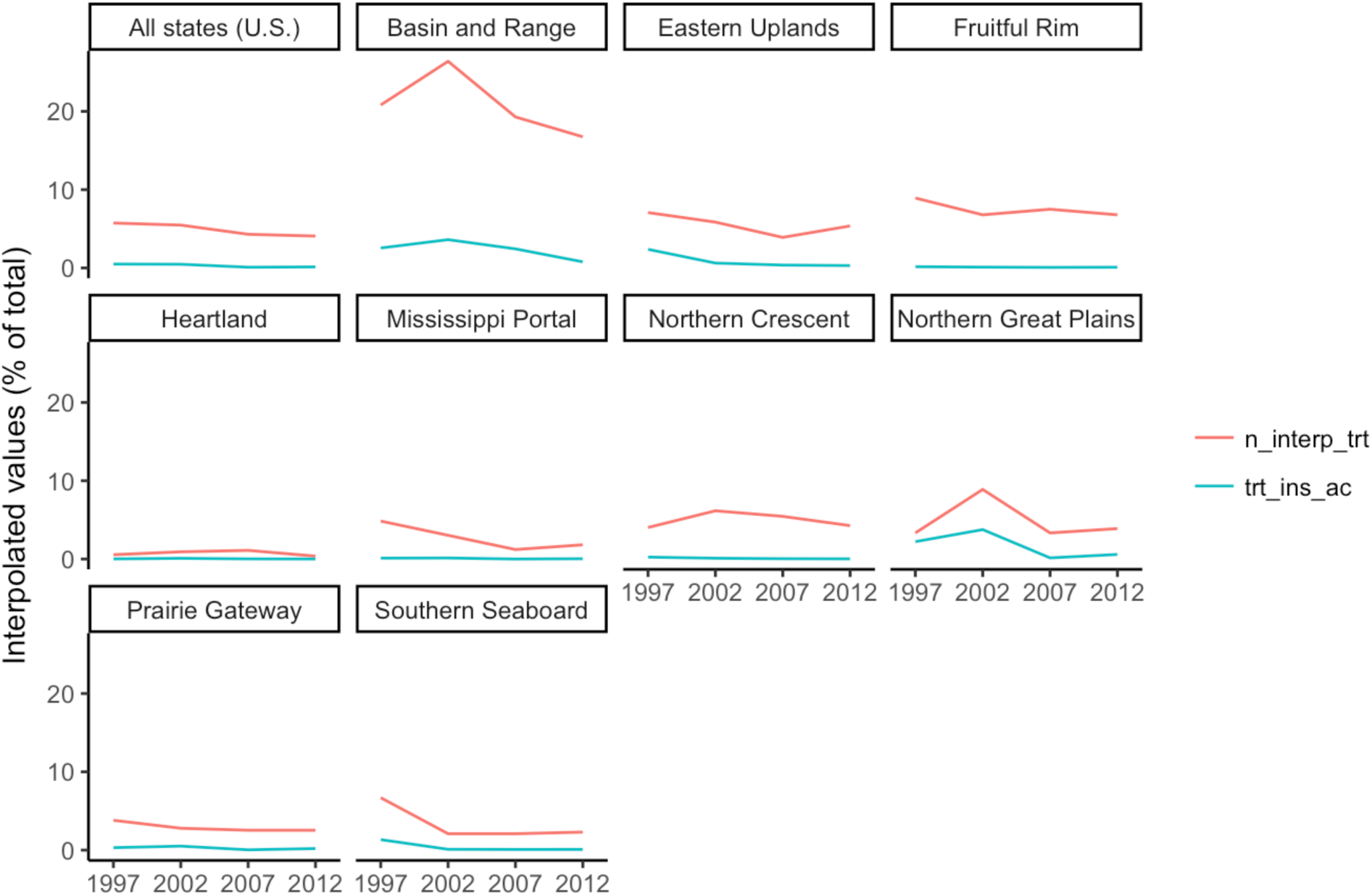
Contribution of counties with interpolated treated area values to total counties (n_interp) and treated cropland (trt_ins_ac) for each of nine agricultural regions and all regions combined.

**Figure S8.**
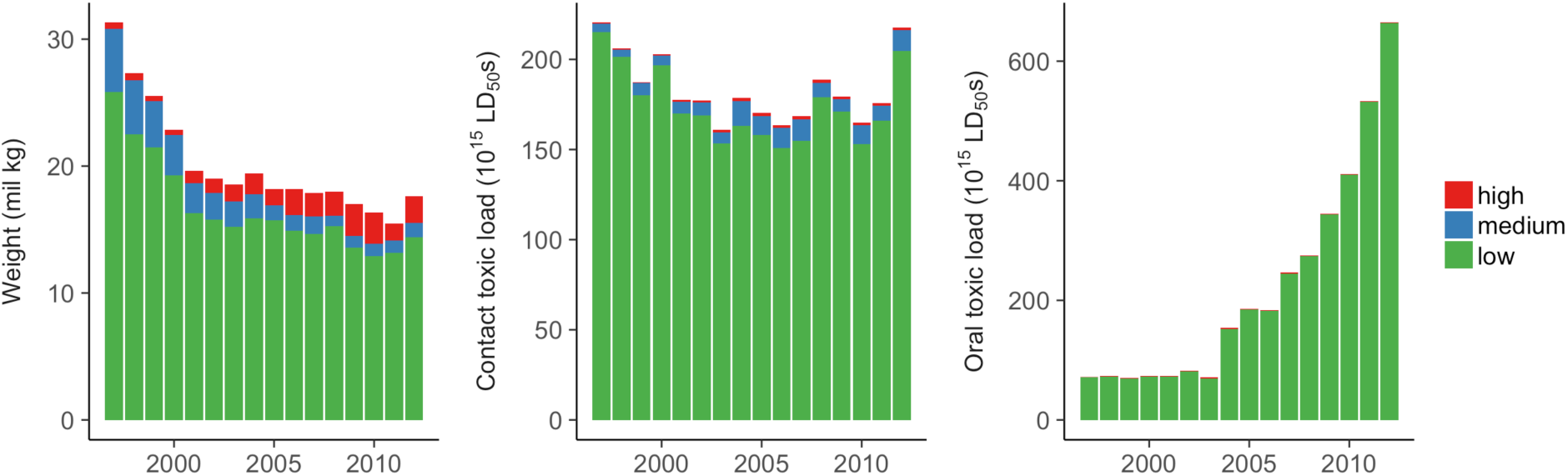
Contribution of LD_50_ values of low, medium, and high uncertainty to national estimates of insect toxic load. LD_50_ values are considered low-uncertainty if they are derived from US or EU regulatory procedures, medium-uncertainty if they are compound-specific but derived from the general scientific literature, and high-uncertainty if they are based on class or insecticide median values.

**Figure S9.**
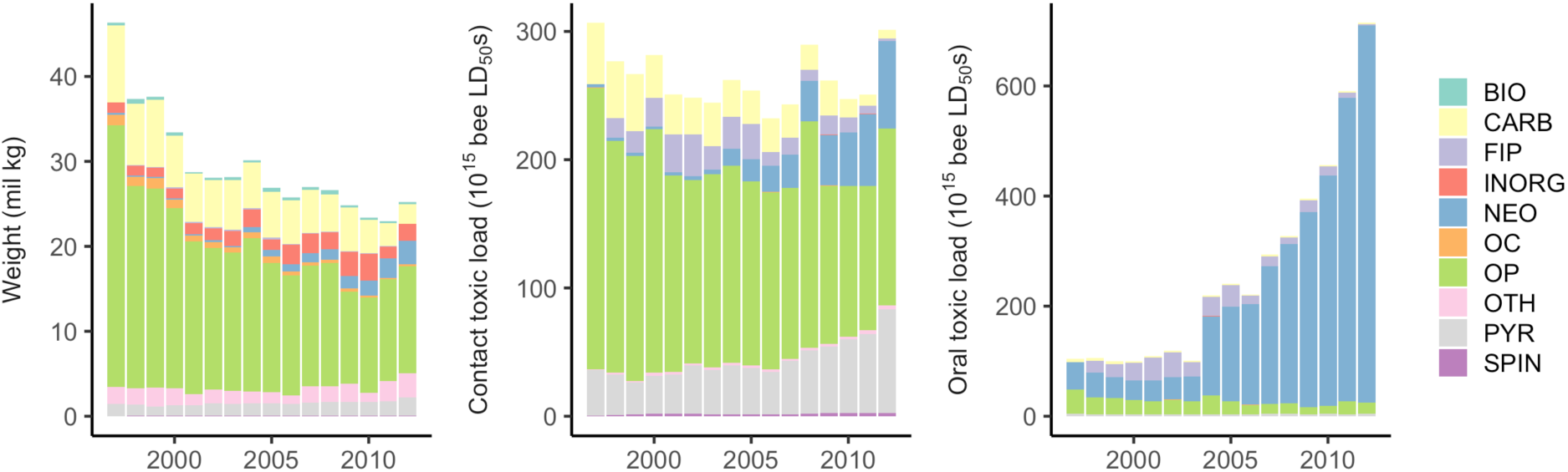
National analysis repeated with the USGS ‘high’ insecticide use estimate.

**Table S1.**
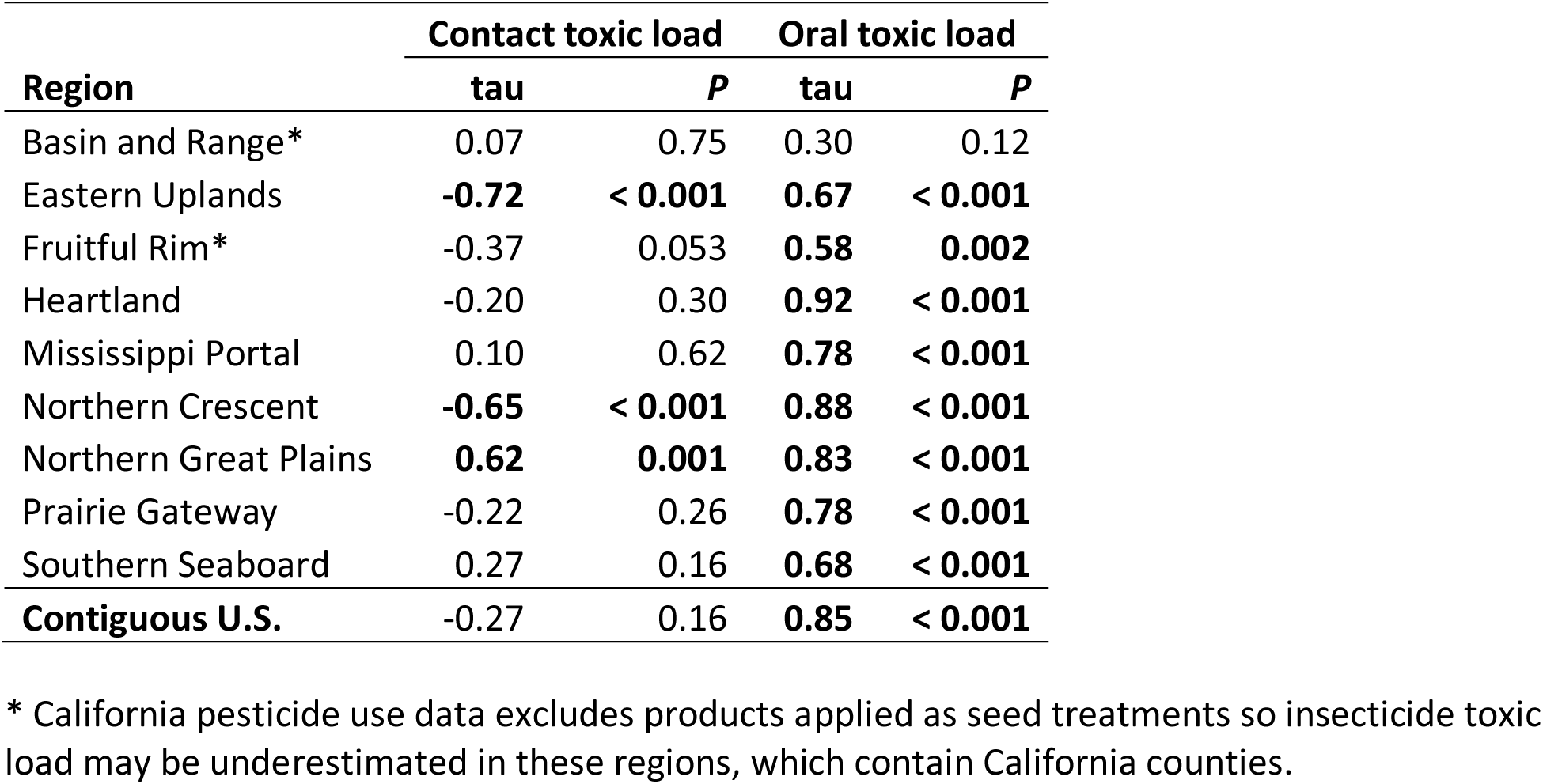
Results from Mann-Kendall trend tests for contact-based and oral-based insect toxic load, from 1997-2012 for nine US agricultural production regions. Tests were considered significant at *P* < 0.05.

**Table S2.**
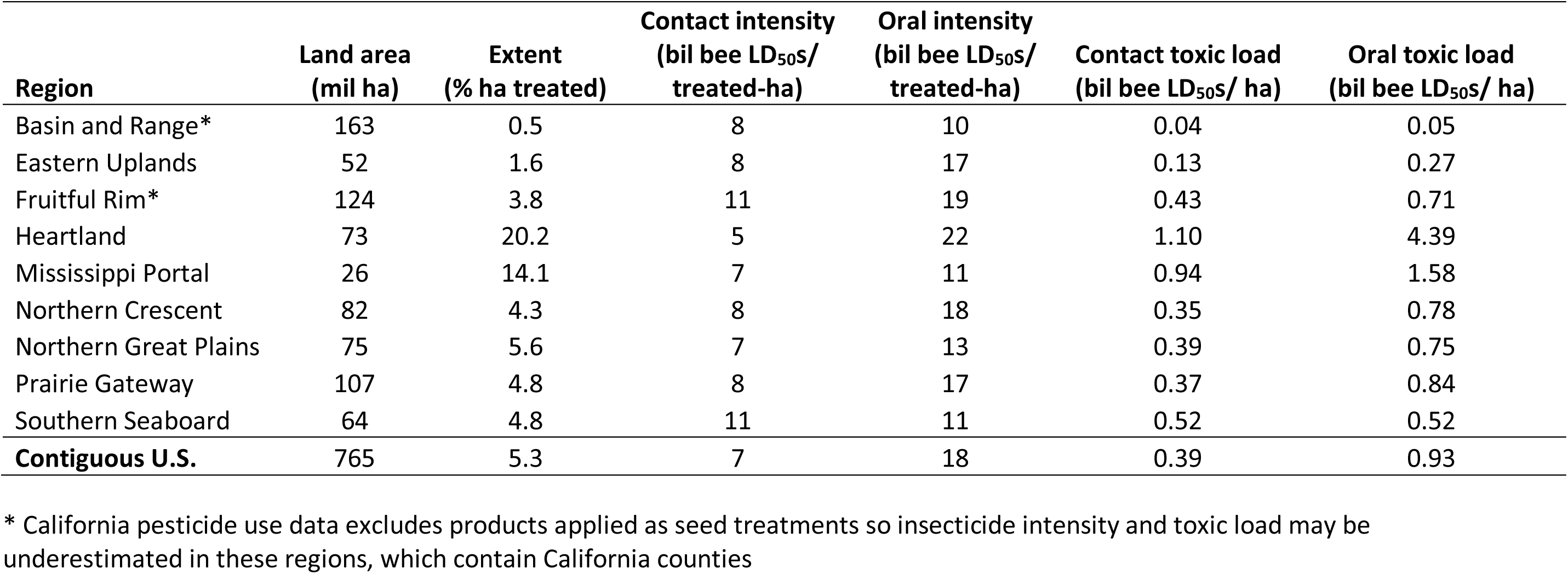
Estimates of 2012 insect toxic load and its contributors for agricultural production regions of the U.S. associated with insecticide use, based on the less-conservative ‘high’ estimate from the USGS National Pesticide Synthesis Project.

