## Supplemental material for "Rising insecticide potency outweighs falling application rate to make US farmland increasingly hazardous to insects"

**SUPPLEMENTAL FIGURES**

**Figure S1.** Honey bee $LD_{50}$ values by insecticide class. Key: FIP = fipronil, NEO = neonicotinoid, SPIN = spinosad, PYR = pyrethroid, CARB = carbamate, OP = organophosphate, OC = organochlorine, INORG = inorganic, BIO = biological, and OTH = other.

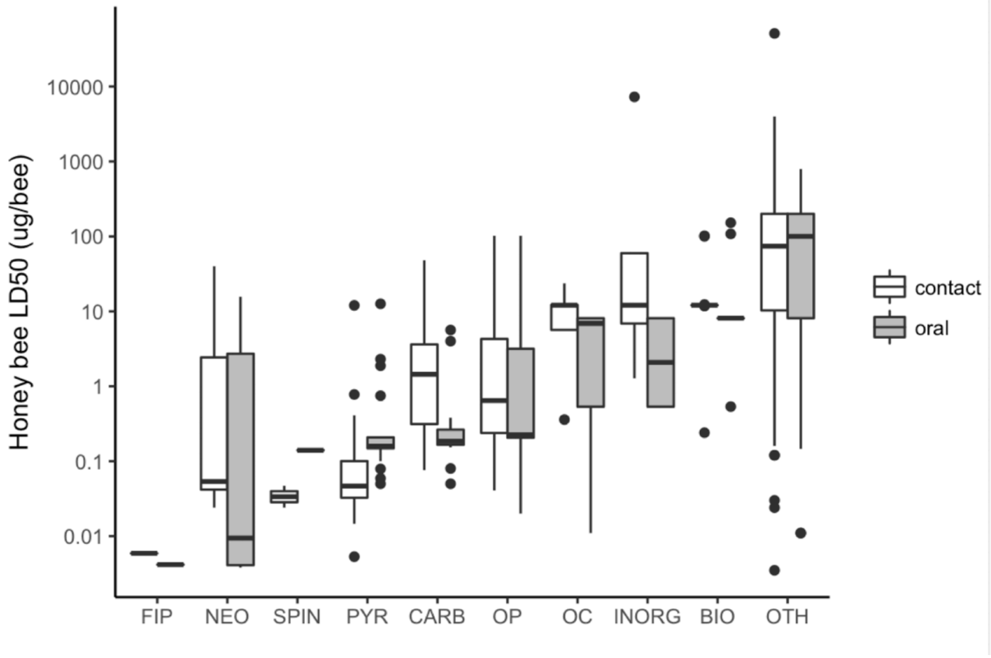

**Figure S2.** Composition of cropland by region in 1997 and 2012, based on crop area data from the U.S. Census of Agriculture.

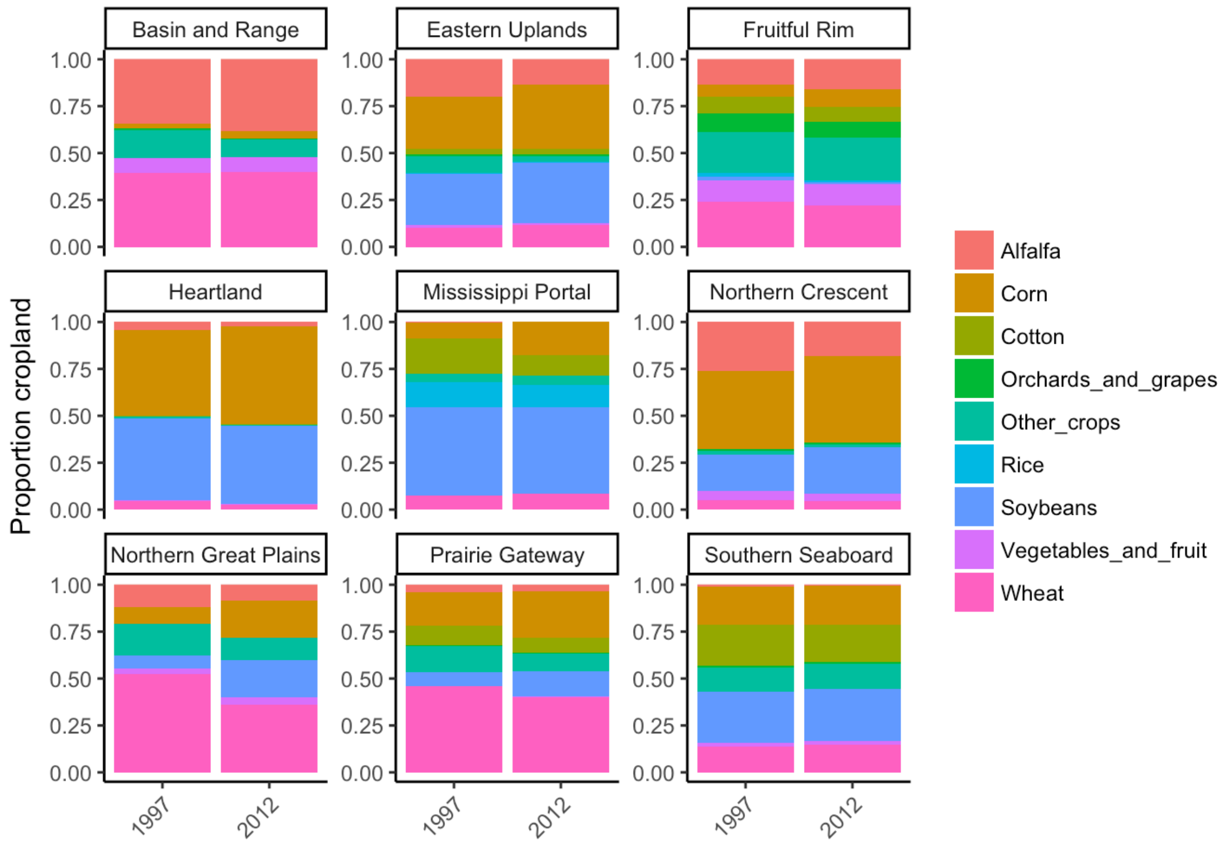

**Figure S3.** Change in the drivers of insect toxic load from 1997 to 2012 for all agricultural production regions, based on the USGS low (left) and high (right) pesticide use estimates. Fold-change is calculated as a response ratio: $Value_{2012}/Value_{1997}$, so that a value of one represents no change, two represents a doubling, one half represents a decline of 50% (values are presented on a log scale).

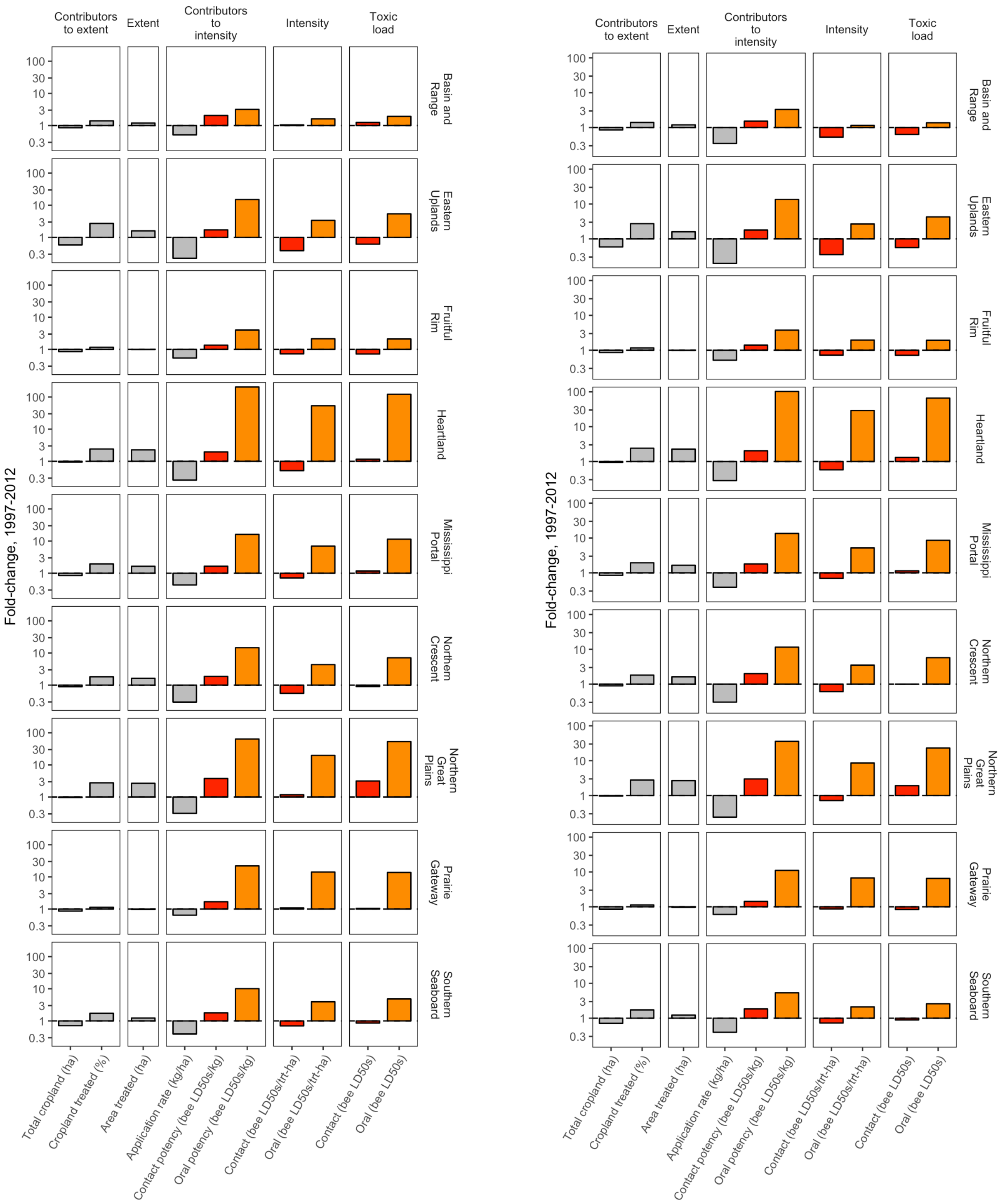

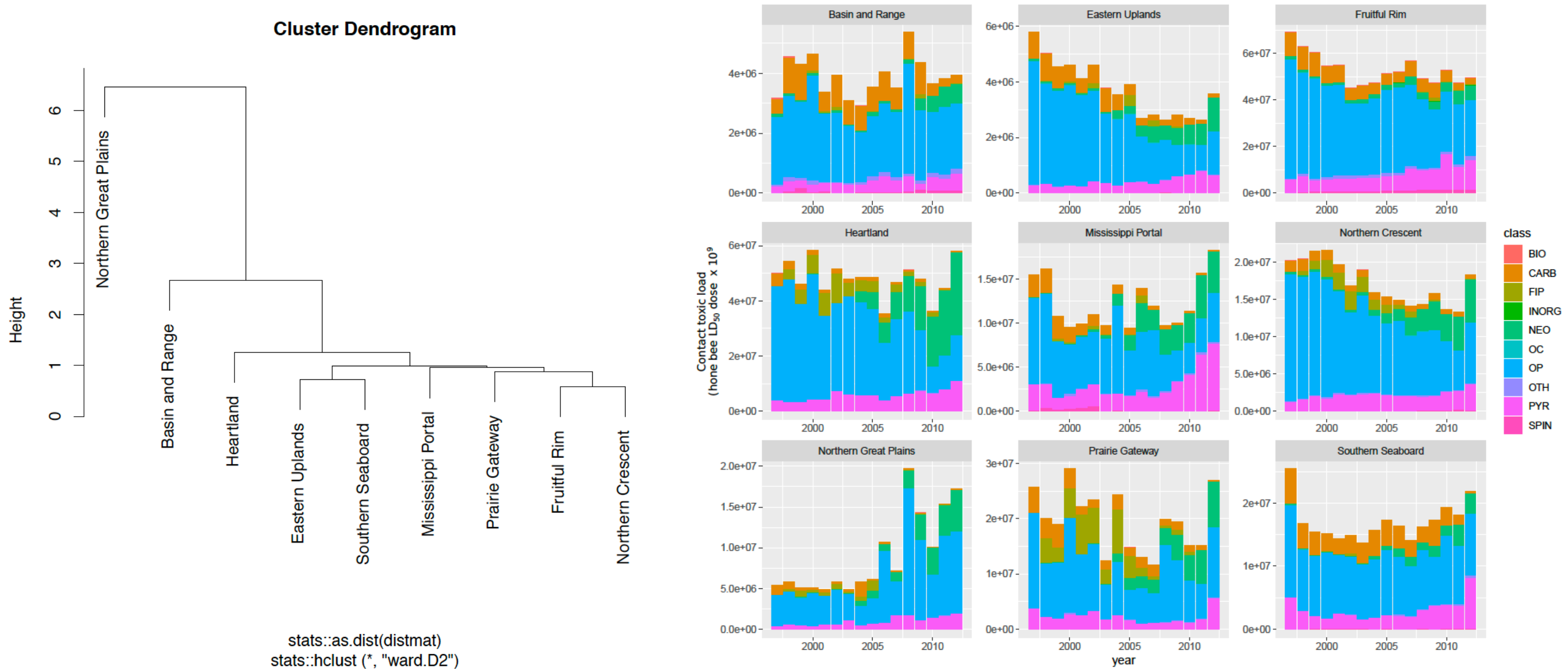


**Figure S4. Contact toxic load by region and chemical class.** Toxic load time series were constructed for each of the nine USDA Farm Resource Regions. Hierarchical clustering (left) grouped regions with similar patterns of toxic load using a Euclidean distance matrix and Ward's linkage method. The y-axis distance between two nodes and their nearest common node is inversely proportional to similarity. Hence, the majority of variation among regions is captured in the first split that separates the Northern Great Plains from the other eight regions. The contribution of different chemical classes to the overall toxic load pattern in each region is depicted with stacked bar plots (right).

**Figure S5.** Contribution of counties with interpolated insecticide values to total counties (n_interp), mass applied (kg_low), contact toxic load (ct_tox), and oral toxic load (or_tox) for each of nine agricultural regions and all regions combined.

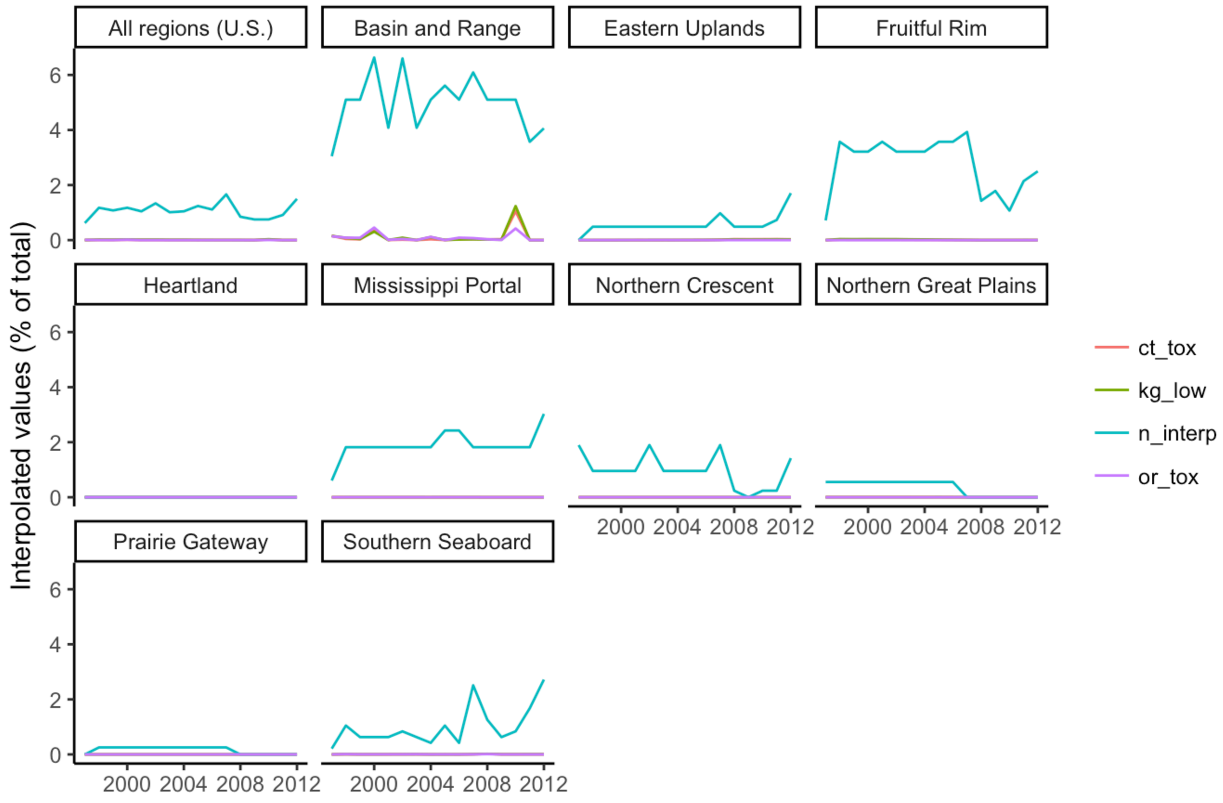

**Figure S6.** Contribution of counties with interpolated cropland area values to total counties (n_interp) and cropland (crop_ac) for each of nine agricultural regions and all regions combined.

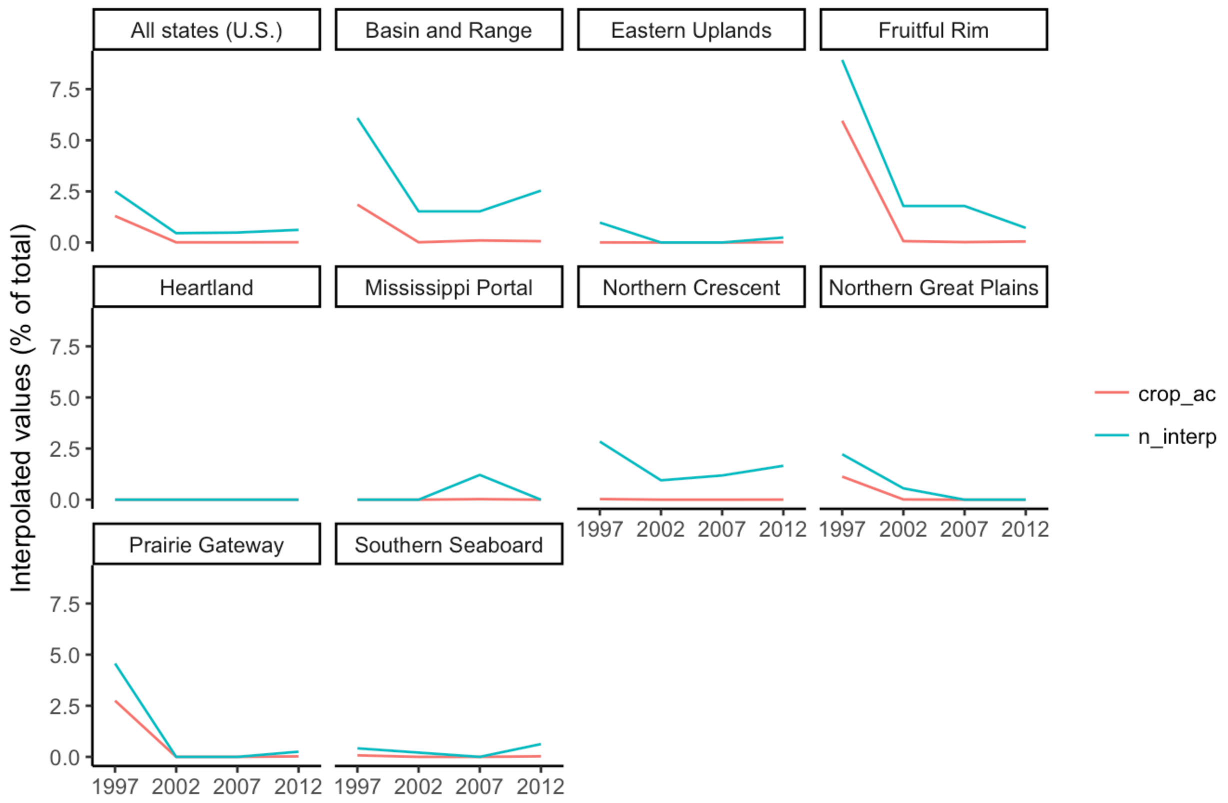

**Figure S7.** Contribution of counties with interpolated treated area values to total counties (n_interp) and treated cropland (trt_ins_ac) for each of nine agricultural regions and all regions combined.

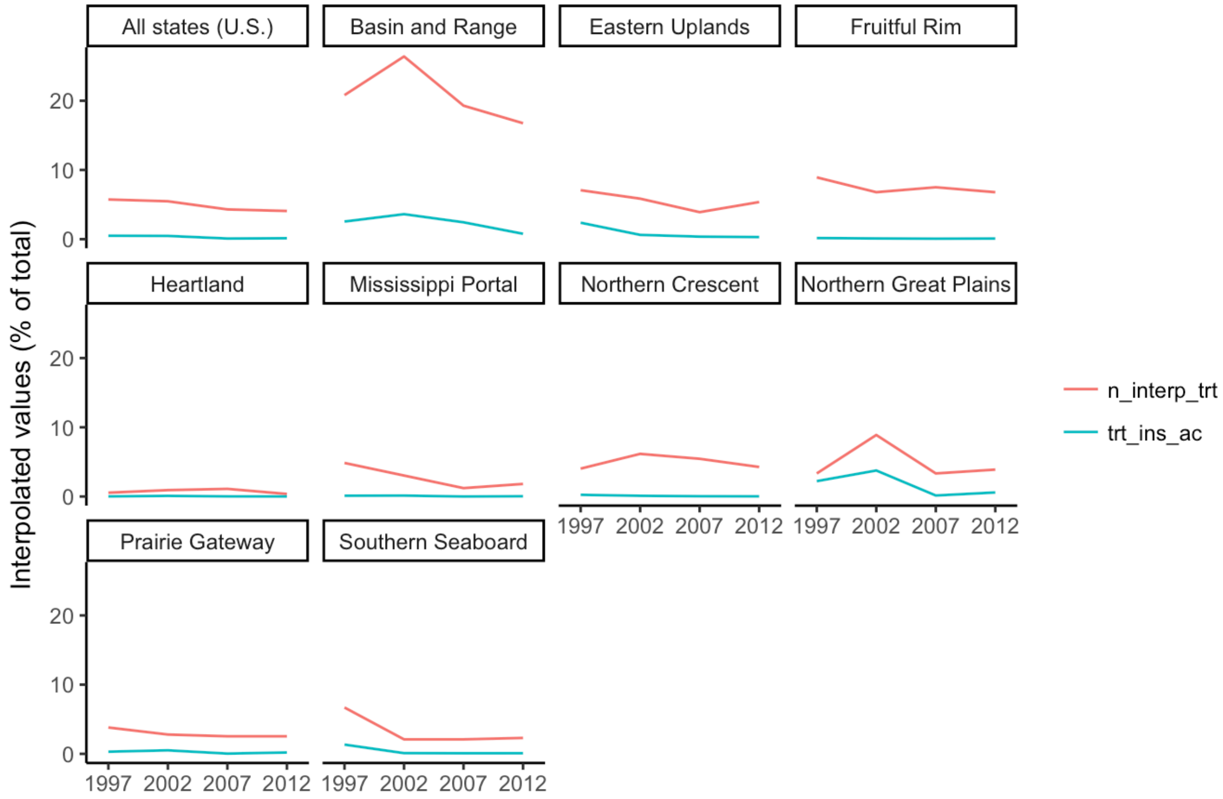

**Figure S8.** Contribution of $LD_{50}$ values of low, medium, and high uncertainty to national estimates of insect toxic load. $LD_{50}$ values are considered low-uncertainty if they are derived from US or EU regulatory procedures, medium-uncertainty if they are compound-specific but derived from the general scientific literature, and high-uncertainty if they are based on class or insecticide median values.

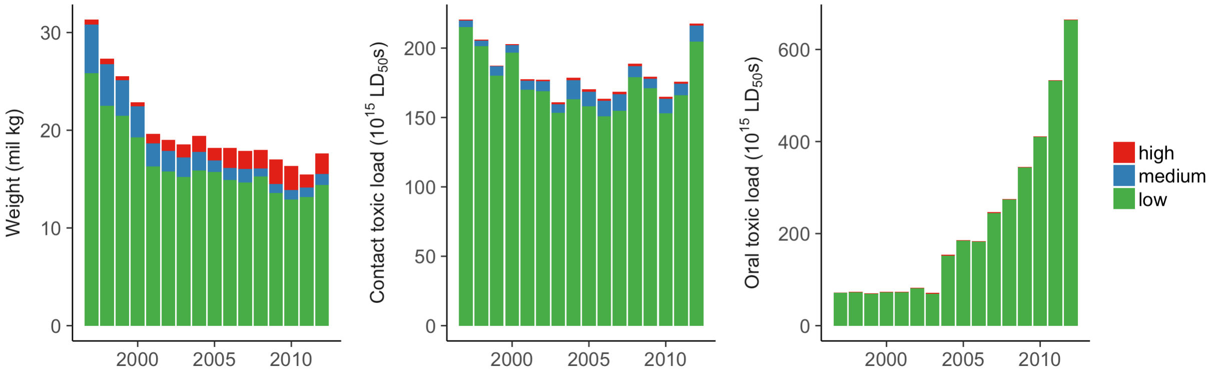

**Figure S9.** National analysis repeated with the USGS 'high' insecticide use estimate.

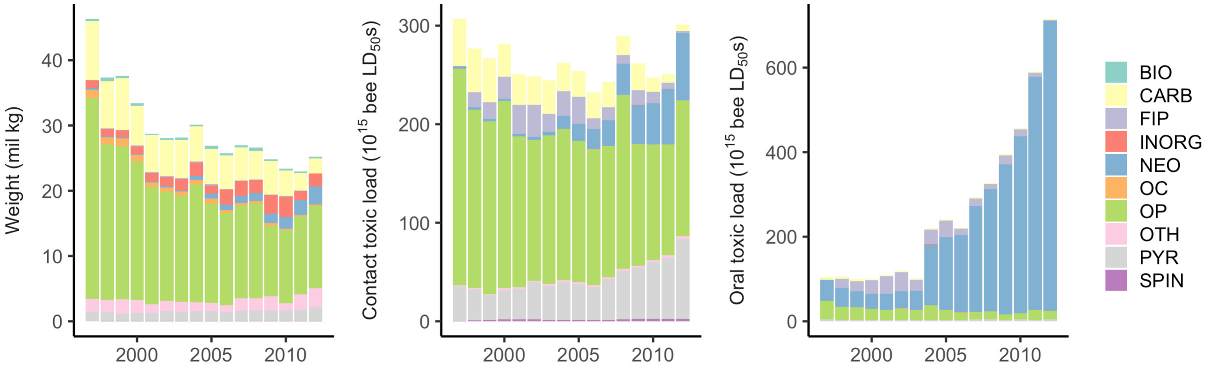

**Table S1.** Results from Mann-Kendall trend tests for contact-based and oral-based insect toxic load, from 1997-2012 for nine US agricultural production regions. Tests were considered significant at $P < 0.05$.

| | Contact toxic load | | Oral toxic load | |
|---|---|---|---|---|
| **Region** | **tau** | ***P*** | **tau** | ***P*** |
| Basin and Range* | 0.07 | 0.75 | 0.30 | 0.12 |
| Eastern Uplands | **-0.72** | **< 0.001** | **0.67** | **< 0.001** |
| Fruitful Rim* | -0.37 | 0.053 | **0.58** | **0.002** |
| Heartland | -0.20 | 0.30 | **0.92** | **< 0.001** |
| Mississippi Portal | 0.10 | 0.62 | **0.78** | **< 0.001** |
| Northern Crescent | **-0.65** | **< 0.001** | **0.88** | **< 0.001** |
| Northern Great Plains | **0.62** | **0.001** | **0.83** | **< 0.001** |
| Prairie Gateway | -0.22 | 0.26 | **0.78** | **< 0.001** |
| Southern Seaboard | 0.27 | 0.16 | **0.68** | **< 0.001** |
| **Contiguous U.S.** | -0.27 | 0.16 | **0.85** | **< 0.001** |

* California pesticide use data excludes products applied as seed treatments so insecticide toxic load may be underestimated in these regions, which contain California counties.

**Table S2.** Estimates of 2012 insect toxic load and its contributors for agricultural production regions of the U.S. associated with insecticide use, based on the less-conservative ‘high’ estimate from the USGS National Pesticide Synthesis Project.

| Region | Land area (mil ha) | Extent (% ha treated) | Contact intensity (bil bee $LD_{50}$s/ treated-ha) | Oral intensity (bil bee $LD_{50}$s/ treated-ha) | Contact toxic load (bil bee $LD_{50}$s/ ha) | Oral toxic load (bil bee $LD_{50}$s/ ha) |
|---|---|---|---|---|---|---|
| Basin and Range* | 163 | 0.5 | 8 | 10 | 0.04 | 0.05 |
| Eastern Uplands | 52 | 1.6 | 8 | 17 | 0.13 | 0.27 |
| Fruitful Rim* | 124 | 3.8 | 11 | 19 | 0.43 | 0.71 |
| Heartland | 73 | 20.2 | 5 | 22 | 1.10 | 4.39 |
| Mississippi Portal | 26 | 14.1 | 7 | 11 | 0.94 | 1.58 |
| Northern Crescent | 82 | 4.3 | 8 | 18 | 0.35 | 0.78 |
| Northern Great Plains | 75 | 5.6 | 7 | 13 | 0.39 | 0.75 |
| Prairie Gateway | 107 | 4.8 | 8 | 17 | 0.37 | 0.84 |
| Southern Seaboard | 64 | 4.8 | 11 | 11 | 0.52 | 0.52 |
| **Contiguous U.S.** | 765 | 5.3 | 7 | 18 | 0.39 | 0.93 |

* California pesticide use data excludes products applied as seed treatments so insecticide intensity and toxic load may be underestimated in these regions, which contain California counties
